# SMARCAD1 Regulates R-Loops at Active Replication Forks Linked to Cancer Mutation Hotspots

**DOI:** 10.1101/2024.09.13.612941

**Authors:** Sidrit Uruci, Nicole M. Hoitsma, María E. Solér-Oliva, Aleix Bayona-Feliu, Vincent Gaggioli, María L. García-Rubio, Calvin S.Y. Lo, Collin Bakker, Jessica Marinello, Eleni Maria Manolika, Giovanni Capranico, Martijn S. Luijsterburg, Karolin Luger, Andrés Aguilera, Nitika Taneja

**Author notes:** shared correspondence. These authors contributed equally to this work (Nicole M. Hoitsma, María E. Solér-Oliva, Aleix Bayona-Feliu).

## Abstract

DNA replication often encounters obstacles like the stalled transcription machinery and R-loops. While ribonucleases and DNA-RNA helicases can resolve these structures, the role of chromatin remodelers remains understudied. Through a series of *in vitro* and *in vivo* experiments, we show that the chromatin remodeler SMARCAD1, which associates with active replication forks, is crucial for resolving nearby R-loops to maintain fork stability. SMARCAD1 directly binds R-loops via its ATPase domain and associates with the replisome through its N-terminus region. Both interactions are critical for resolving R-loops within cells. Genome-wide assays reveal that cells expressing mutant SMARCAD1 accumulate significantly more R-loops than wild-type cells, particularly in regions distinct from known fork blockage-prone sites. These R-loop-enriched regions in SMARCAD1 mutants also exhibit increased mutagenesis in germline tumors, suggesting they are mutation hotspots in cancer. Therefore, SMARCAD1 acts as an R-loop sensor and resolvase at actively progressing forks, maintaining genome stability and preventing tumorigenesis.

## INTRODUCTION

Transcription is a fundamental process that can become an obstacle to replication fork progression, resulting in transcription-replication conflicts (TRCs)^1–6^. These conflicts are often associated with the formation of R-loops, structures composed of a DNA-RNA hybrid and a displaced section of single-stranded DNA (ssDNA). The genesis of R-loops can be either scheduled or unscheduled. While scheduled and transient R-loops are linked to physiological functions such as class-switch recombination^7^, unscheduled and persistent R-loops are widely associated with replication stress that causes genome instability, a hallmark of cancer^8–12^.

In response to R-loop accumulation, cells have evolved intricate preventive mechanisms involving factors assembling onto nascent RNA or relieving topological stress^13^. Furthermore, cells also have means to remove R-loops either via nucleases (such as RNaseH1^14,15^ and DICER^16^) or DNA–RNA helicases (like SETX^17^, AQR^18^, UAP56^19^ and others). Finally, to successfully counteract their negative impact, the damage caused by R-loops is repaired^20,21^. While considerable focus has been devoted to delineating pathways preventing R-loop accumulation, only recently has attention shifted to the role of the chromatin landscape in R-loop processing^22–28^. Chromatin reorganization has been linked to the proper resolution of R-loops by altering nucleosome positioning, thereby improving the accessibility of R-loops to resolvases^29^. However, a key unresolved question remains: how do chromatin remodelers and modifiers, associated with specific states of replication forks, distinguish between the persistent and detrimental accumulation of DNA-RNA hybrids, which pose obstacles ahead of the fork, and the transient, scheduled hybrids linked to normal replisome function? This uncertainty has led to the hypothesis that there could be distinct resolving activities associated with both unperturbed and stressed replication forks^30^. In this scenario, SWI/SNF chromatin remodelers, such as BRG1 may mediate R-loop resolution at blocked replication forks, alleviating the conflict and mediating repair at the stressed site to facilitate their restart^22^. In unperturbed conditions, it is hypothesized that chromatin-associated factors at active replication forks can sense and interact with TRCs or R-loops ahead of the forks, processing these conflicts through direct interaction without stalling the fork or inducing a DNA damage response (DDR)^30–33^.

SMARCAD1, a protein with a DEAD/H box helicase domain, is part of the highly conserved ATP-dependent INO80 family of chromatin remodelers^34–37^. SMARCAD1 and its homolog in budding yeast (Fun 30) are known to play roles in various biological processes, including transcription^38,39^, heterochromatin maintenance^40,41^, and DNA repair^34–36^. While the fission yeast Fun30 homolog, Fft3, is known to interact with DNA-RNA hybrids, the mechanistic implications of this interaction remain unclear^41^. Recent research from our lab has uncovered a novel role for human SMARCAD1 in fine tuning PCNA levels at replication forks, ensuring their stability to maintain genome integrity^42^.

In this study, we describe a new role for SMARCAD1 in processing R-loops in proximity of active replication forks. Utilizing functional mutants of human SMARCAD1, we captured high levels of TRCs and R-loops in replication-associated defective and nucleosome remodeling defective-mutant cells. Through *in vitro* biochemical assays, single molecule atomic force microscopy, *in vivo* artificial tethering, and genetic studies, we demonstrate that SMARCAD1 directly binds to R-loops and facilitates their processing in the vicinity of active replication forks. The regions accumulating R-loops in SMARCAD1-mutant cell lines are distinct from those previously identified as prone to replication fork breakage^22^. Additionally, regions with R-loop accumulation in SMARCAD1 mutant cells exhibit increased cancer-associated mutagenesis. Our study unveils SMARCAD1 as an autonomous remodeler capable of directly binding the R-loops, thus facilitating their processing, and TRCs ahead of replication forks, emerging as a key player in maintaining genome stability.

## RESULTS

### SMARCAD1 is a potential R-loop-binding protein associated with active replication forks

Chromatin factors have emerged as critical players in maintaining R-loop homeostasis and mitigating associated genome instability^43–45^. While recent studies have focused on the role of these chromatin factors in processing R-loops, the specific states of replication forks at which they act remain largely unexplored. This is particularly important given the dual nature of R-loops—persistent versus transient—and the possibility that distinct pathways may process them differently, depending on whether they accumulate at unstressed and progressing forks or stressed, slowed down, stalled forks. Understanding these dynamics is crucial for fully grasping their role in genome stability.

We hypothesize that chromatin remodelers interacting with actively progressing replication fork may sense TRCs and R-loops, and process them through direct physical interactions. This mechanism would prevent fork stalling and the activation of the DDR. To test this hypothesis and to identify chromatin modifiers that interact with R-loops when associated with the active fork, we mined a previous R-loop proximity proteomics screen^46^. When reanalyzing this dataset, we specifically focused on highly and significantly enriched (change > 2-fold and p-value < 0.01) chromatin modifiers and remodelers. We identified thirty-one factors that are enriched at R-loops (Extended Data Fig. 1a and Table 1). We then cross-referenced these results with our previous isolation of proteins on nascent DNA (iPOND) coupled to stable isotope labeling with amino acids in cell culture (SILAC)–based quantitative mass spectrometry dataset^42^. This analysis revealed that ∼70% of these factors are generally enriched at replication forks, with only a few having a preference for the active fork (Extended Data Fig. 1b and Table 1). One of the most enriched chromatin remodelers identified through this analysis is SMARCAD1 (SWI/SNF-related matrix-associated actin-dependent regulator of chromatin subfamily A containing DEAD/H box 1) (Extended Data Fig. 1a, b), which associates with unperturbed/active replication forks and has been previously predicted to bind R-loops through bioinformatic analysis^47^.

**Table 1.**
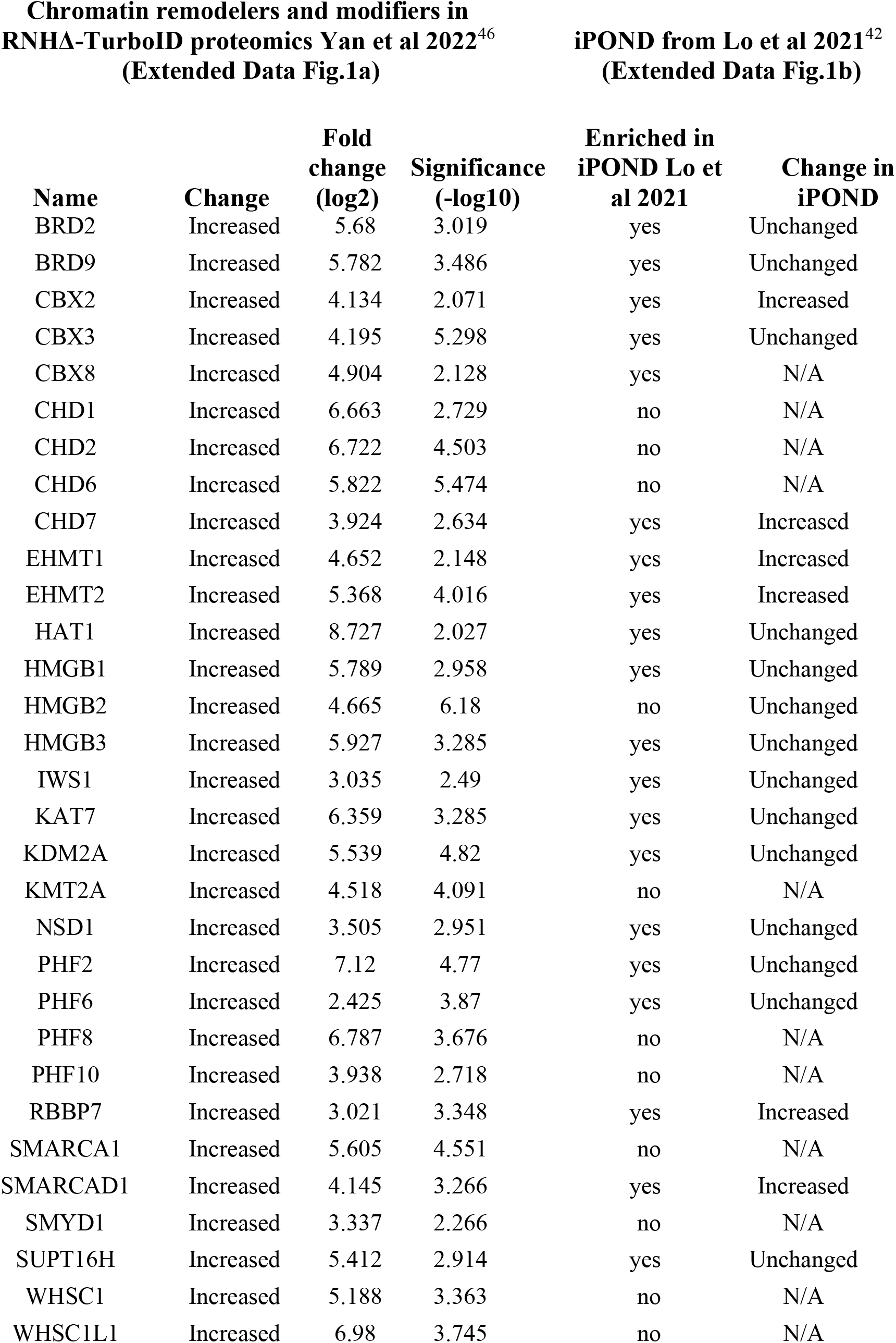

SMARCAD1 has been characterized for its roles in DNA damage repair^48^, and in heterochromatin maintenance^40,49^, but also in a novel pathway stabilizing the active replication fork by maintaining PCNA homeostasis^42^. However, it remains unclear whether SMARCAD1, in its association with active forks, can sense conflicts and R-loops to facilitate their resolution without causing fork stalling.

### SMARCAD1 binds DNA-RNA hybrids and R-loop structures *in vitro*

To validate the possible interaction of SMARCAD1 with R-loops, we generated an “R-loop-like” substrate (referred to as R-loopshort) that contains a bubble of single-stranded DNA (ssDNA, 31 nt) flanked by two arms of double-stranded DNA (dsDNA, 30 bp), with an RNA (25 nt) that anneals to form a DNA-RNA hybrid in the bubble (Extended Data Fig. 2a). The R-loopshort substrate was designed to have minimal flanking dsDNA and a defined loop region, as opposed to transcriptionally generated R-loops which are heterogeneous in size (described below). The structure of the annealed R-loopshort substrate was visualized via Atomic Force Microscopy (AFM), where height analysis confirms the DNA-RNA hybrid and dsDNA flanking the hybrid, as well as the opposing ssDNA (Extended Data Fig. 2b, c). These features were also confirmed by mobility in native gels and restriction digest, which utilize the fluorescein (FAM) label on the RNA oligonucleotide to confirm its annealing within the ssDNA bubble, as shown previously^50^ (Extended Data Fig. 2d). The fluorescent RNA oligo was also annealed to a complementary DNA sequence to form a DNA-RNA hybrid substrate, which was confirmed by native gel (Extended Data Fig. 2a, e). We then employed *in vitro* biochemical analysis to study the binding of SMARCAD1 to these substrates, via electrophoretic mobility shift assay (EMSA). After incubating each substrate with purified SMARCAD1, we observed reduced mobility of the complex compared to the free nucleic acid substrates, confirming SMARCAD1 interaction (Fig. 1a). To quantify SMARCAD1 binding, we monitored fluorescence polarization of the FAM labeled substrates at increasing concentrations of SMARCAD1 (Fig. 1b). This data reveals that SMARCAD1 binds DNA-RNA hybrid and R-loopshort substrates with a high affinity that is comparable to dsDNAshort with a <2-fold difference in EC50 values (Fig. 1c). While the highest affinity is observed for the R-loopshort, we cannot distinguish SMARCAD1 binding different portions of the substrate (*i.e.* DNA-RNA hybrid, R-loop, dsDNA, ssDNA). However, the binding affinity for ssDNA is >4-fold reduced compared to the hybrid and R-loop, suggesting that SMARCAD1 may preferentially interact with the DNA-RNA hybrid (Fig. 1c).

**Figure 1.**
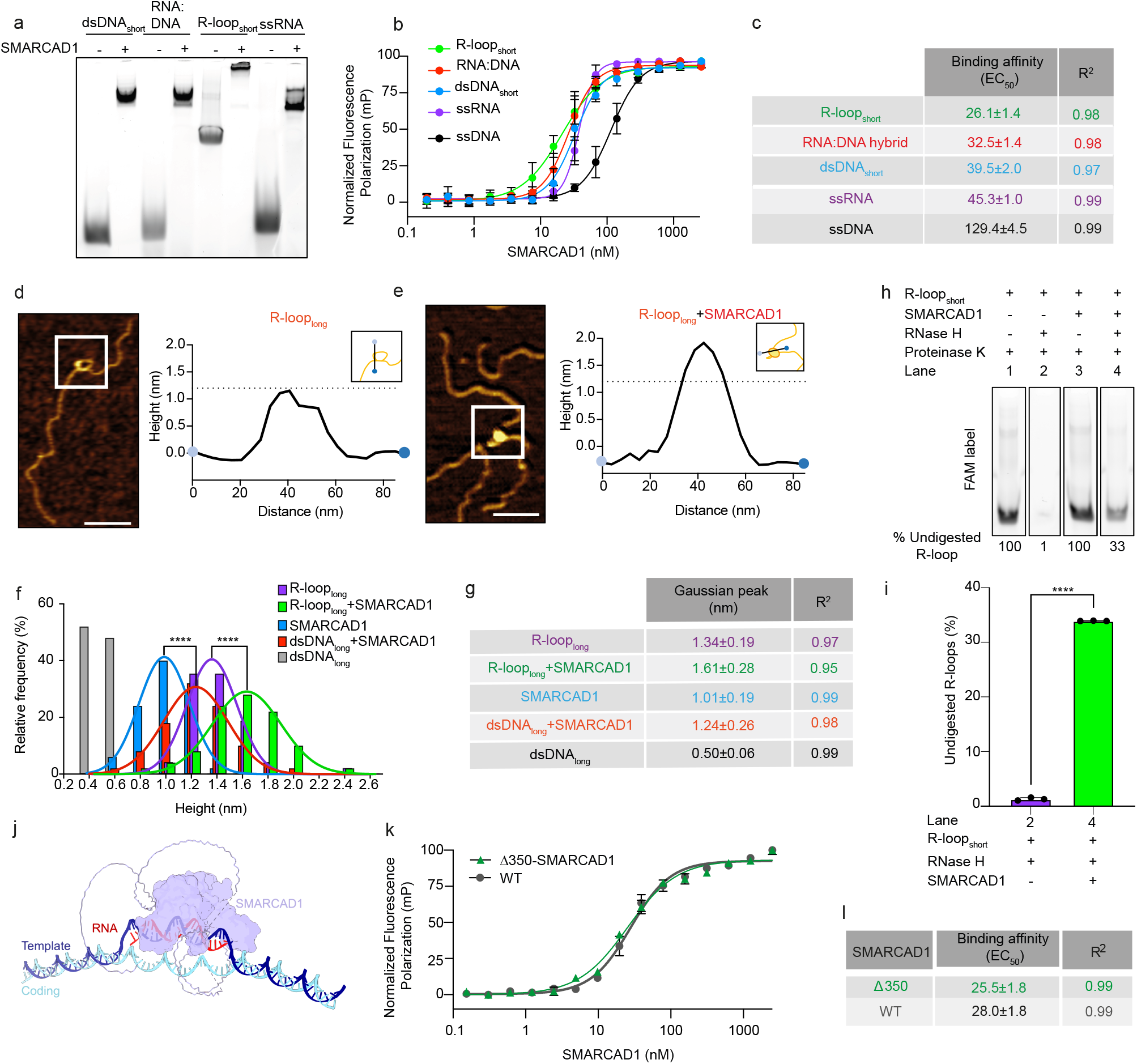
Chromatin remodeler SMARCAD1 binds *in vitro* R-loops. **a.** Each substrate in the presence or absence of excess SMARCAD1, analyzed by EMSA on a 5% 59:1 acrylamide gel, visualized by FAM fluorescence. Uncropped gel available in Source Data. **b.** Fluorescence Polarization assay of SMARCAD1 (0 to 2500 nM) binding FAM labeled substrates. Data points shown are the mean ± standard deviation of three independent replicates. Numerical data available in Source Data. **c.** EC50 binding affinity of the indicated substrates, mean ± standard error and R^2^ of curve fit shown in **b**. **d.** Left: Representative image of AFM analysis for R-looplong. Right: AFM height profile through the point of maximum height as depicted in the graphical inset. Dashed line at 1.2 nm height shown for reference. Scale bar, 100 nm. **e.** Left: Representative image of AFM analysis for R-looplong + SMARCAD1. Right: AFM height profile through the point of maximum height as depicted in the graphical inset. Dashed line at 1.2 nm height shown for reference. Scale bar, 100 nm. **f.** Histogram and gaussian fitting of AFM height analysis (n=50). Gaussian fit for dsDNAlong is not shown for clarity. **g.** Observed height corresponding to the center of each peak, shown in **f**, as mean ± standard deviation and R^2^ of Gaussian fit. **h.** Representative protection assay gel visualized for FAM label on RNA oligonucleotide. Samples were treated with Proteinase K to digest any protein after the reaction, therefore only the unbound R-loop substrate is visualized in the gel. Below gel is quantification of RNA bands in the indicated conditions. Lane 1-4 are taken from Extended Data Fig. 2i. **i.** Protection assay quantification RNA band (FAM label in **h**) as percent of respective undigested band (n=3, SD shown as error bars). **j.** Cartoon representation of AlphaFold3 prediction of full-length WT SMARCAD1 (purple, N-terminal region shown as unstructured loop, C-terminal region shown as transparent surface) and R-loopshort substrate. Overall, the Alphafold3 prediction was of low confidence with ipTM = 0.29 and pTM = 0.53. However, the chain pair iPTM for SMARCAD1 and the RNA is 0.58, with this higher confidence also represented in the predicted aligned error. **k.** Fluorescence Polarization assay of WT and Δ350-SMARCAD1 (0 to 2500 nM) binding the R-loopshort substrate. Data shown are the mean ± standard deviation of technical duplicate. Numerical data available in Source Data. **l.** EC50 binding affinity of the indicated protein, mean ± standard error and R^2^ of curve fit shown in **k**.

An R-loop, by definition, is flanked by dsDNA regions. Since bulk binding analysis cannot preclude SMARCAD1 binding these dsDNA regions, we sought to analyze this interaction at the single-molecule level via AFM. We utilized a previously established system to generate R-loops suitable for AFM analysis via *in vitro* transcription (IVT) of a plasmid containing the mAIRN gene^51^ (Extended Data Fig. 2f). IVT of the mAIRN gene has been shown to form R-loops positioned 38-42% from the closest DNA end of a linear 2745 bp DNA fragment (here, referred to as R-looplong), which has been used to directly visualize the binding of PARP1 via AFM^51^. In agreement with previous reports, we were able to visualize R-loops in three different shapes; “loops,” “spurs,” and “blobs” at ∼40% contour length (Extended Data Fig. 2g)^51–53^. For AFM height analysis, the maximum height was extracted from manually identified particles (Fig. 1d, e). As previously observed^51^, R-looplong particles display a greater height compared to dsDNAlong (Fig. 1f, g). To investigate its binding, SMARCAD1 was incubated with dsDNAlong or R-looplong before deposition on mica for AFM analysis. Imaging of dsDNAlong with SMARCAD1 reveals identifiable SMARCAD1 particles bound to dsDNAlong (Extended Data Fig. 2h). As expected, SMARCAD1 bound to dsDNAlong has a significantly greater height than dsDNAlong or SMARCAD1 alone (Fig. 1f, g). Due to the complex heterogeneous shapes observed for the R-looplong, height analysis is required to infer SMARCAD1 binding to the R-looplong substrate (Fig. 1e-g). Our data show a SMARCAD1-dependent enrichment of R-loops with significantly greater height than R-looplong alone (Fig. 1f, g). While some R-loops are likely unbound, this data suggests that SMARCAD1 binds a population of R-looplong particles to cause the shift observed in the height distribution (Fig. 1f, g).

To further validate its interaction with the R-loop, we tested whether SMARCAD1 binding protects the DNA-RNA hybrid from nuclease cleavage. We used RNaseH1 enzyme, which specifically cleaves the RNA moiety of a DNA-RNA hybrid^54^. Under these conditions, RNaseH1 digests nearly all the R-loopshort substrate, as evidenced by the loss of FAM-labeled RNA in the gel (Fig. 1h). However, upon the addition of SMARCAD1, a significant portion of the R-loopshort substrate remained undigested, suggesting that SMARCAD1 interacts with the hybrid portion of the R-loop substrate. (Fig. 1h, i and Extended Data Fig. 2i). Together, these data suggest that SMARCAD1 directly binds DNA-RNA hybrids and R-loop structures *in vitro*.

### The ATPase domain of SMARCAD1 binds R-loop

Unlike other multi-subunit chromatin remodelers in the INO80 family, SMARCAD1 is a single-unit autonomous remodeler^34^. Based on this, we sought to determine which region of SMARCAD1 interacts with R-loops. For this purpose, we first used the newly released version of AlphaFold3 software, which allows *in silico* prediction of protein folding and interaction with nucleic acid structures^55^. Though the software is still in its beta phase and R-loop structure predictions are of low confidence, this prediction provides a potential model of how SMARCAD1 may interact with an R-loop. As anticipated from the SMARCAD1 sequence (UniProt ID: Q9H4L7), the Alphafold3 prediction shows a mostly unstructured N-terminus and a structured C-terminus containing extended ATPase domains (350-1026). Notably, AlphaFold3 reproducibly positions the C-terminal region of SMARCAD1 on the DNA-RNA hybrid, and not the displaced ssDNA strand or the dsDNA (Fig. 1j). This prediction suggests, albeit at a low confidence, that the catalytic ATPase domains of the C-terminal region of SMARCAD1 directly interact with the DNA-RNA hybrid component of R-loops.

To corroborate this *in silico* prediction, we tested binding of a SMARCAD1 mutant lacking the first 350 amino acids (Δ350-SMARCAD1) via the previously described *in vitro* fluorescence polarization assay. This mutant lacks the flexible N-terminal region while leaving intact the split catalytic domain (ATPase1 and ATPase2) of the C-terminus. This data reveals that Δ350-SMARCAD1 binds the R-loopshort substrate with the same high affinity as SMARCAD1^WT^ (Fig. 1k, l), suggesting that the C-terminus of SMARCAD1 binds directly to R-loops, while the more flexible N-terminus is dispensable for this high-affinity interaction. This is in agreement with the AlphaFold3 prediction.

### Artificial tethering of SMARCAD1 promote R-loop processing *in situ*

After demonstrating the *in vitro* binding of SMARCAD1 to R-loops and DNA-RNA hybrids, we investigated its ability to process these secondary structures *in situ*. To do this, we employed previously described artificial tethering approaches^25,56–58^. We utilized human U2OS 2-6-3 cells containing a 4Mbp heterochromatic region with multiple copies of a 200 repeats tandem array harboring the lactose operator (LacO) sequence^59^. This LacO array serves as a known hotspot for replication stress and R-loop formation^25^. To validate the presence of R-loops at the array locus in our assay, we transfected U2OS 2-6-3 cells with a LacR-encoding plasmid and detected R-loop accumulation at the array locus via immunostaining with the S9.6 antibody, which recognizes the DNA-RNA hybrid of the R-loops^60–62^ (Fig 2a, b). Given that recent studies have suggested a cross-reactivity of this antibody with dsRNA^63,64^, we validated the specificity of the S9.6 antibody for detecting DNA-RNA hybrids before proceeding. We pre-treated cells with RNaseH1 enzyme, which specifically removes the RNA moiety of DNA-RNA hybrids^54^, and RNase III, which degrades dsRNA^65^. We observed a clear reduction in genomic DNA-RNA hybrid levels in cells pre-treated with RNaseH1 enzyme, but not in those treated with RNase III, confirming the specificity of our antibody for DNA-RNA hybrids (Extended Data Fig. 3a, b). Dot-blot analysis using the S9.6 antibody further supported this specificity, as RNaseH1 treatment alone reduced S9.6 intensity, whereas RNase III treatment did not (Extended Data Fig. 3c). These results provided confidence in the ability of our antibody to detect *bona fide* DNA-RNA hybrids.

**Figure 2.**
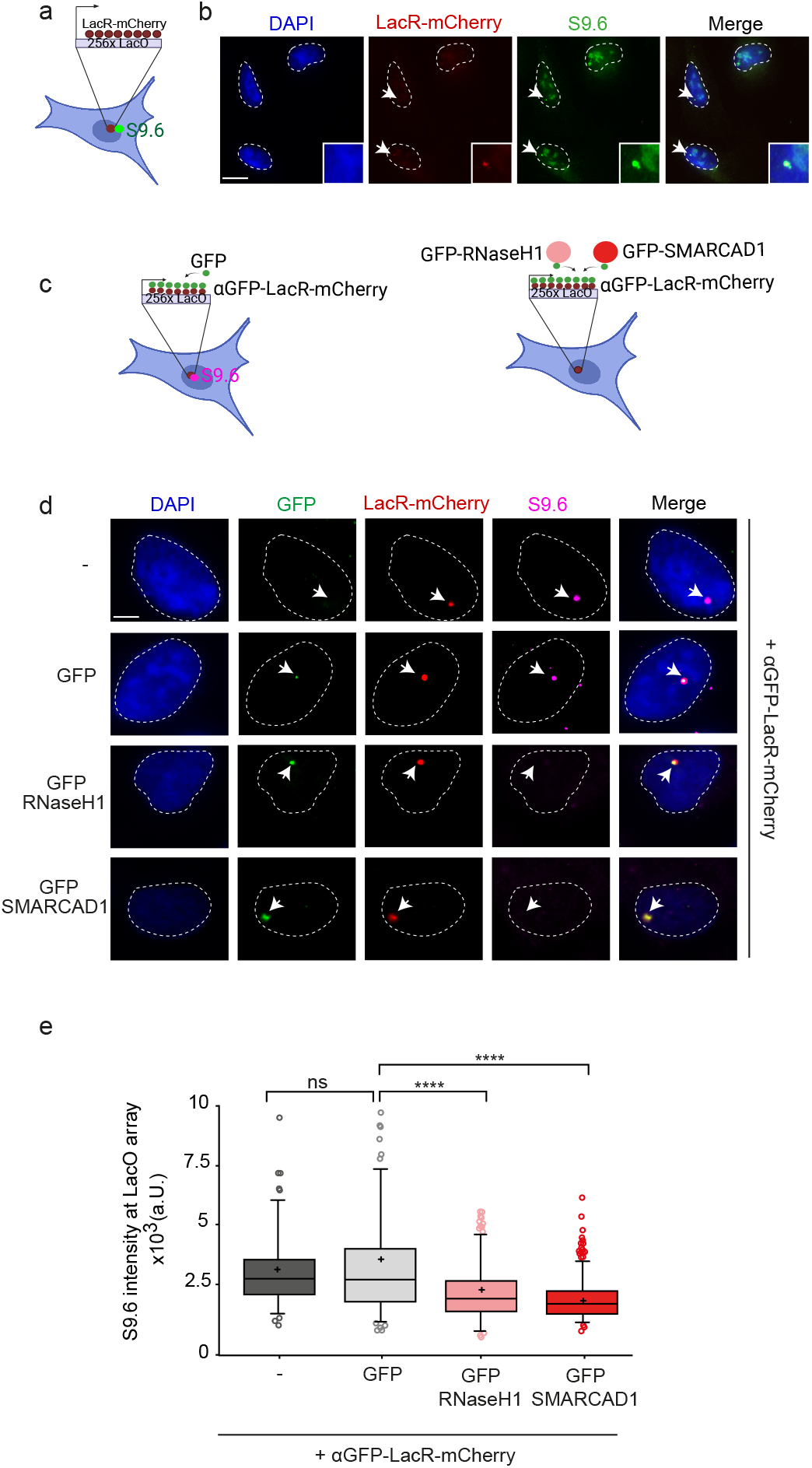
Artificial tethering of SMARCAD1 at LacO locus resolves R-loops *in situ*. **a.** Cartoon of LacO-LacR system for artificial tethering **b.** Representative images of R-loops immunofluorescent labeling using S9.6 antibody at the LacO locus, visualized by artificial tethering of LacR-mCherry. Scale bar, 20 μm. **c.** Cartoon of LacO artificial tethering by αGFP-LacR-mCherry with GFP, GFP-RNaseH1 and GFP-SMARCAD1. **d.** Representative images of R-loops immunofluorescent labeling using S9.6 antibody at the LacO locus, visualized by artificial tethering of αGFP-LacR-mCherry of the specified conditions. Scale bar, 10 μm. **e.** Boxplot representation of the quantification of S9.6 signal at the LacO locus, identified by αGFP-LacR-mCherry from **d**. n(-) = 204 foci, n(GFP) = 242 foci, n(GFP-RNaseH1) = 398 foci, n(GFP-SMARCAD1) = 402 foci from 2 independent replicates. ns = non significant, ****=p≤0.0001, one-way analysis of variance Kruskal-Wallis test, followed by Dunn’s correction for multiple testing. Numerical data available in Source Data.

To target proteins of interest to these sites, we utilized a plasmid encoding LacR-mCherry tagged with a single-domain GFP antibody, resulting in a fusion protein (αGFP-LacR-mCherry)^56^ that efficiently tethered co-expressing GFP-tagged proteins at the LacO tandem array site (Fig. 2c, d). Immunofluorescence analysis using S9.6 in U2OS 2-6-3 cells co-transfected with the fusion protein αGFP-LacR-mCherry along with GFP protein revealed once more the enrichment of R-loops at the LacO array now decorated with GFP while showing no R-loop processing, as expected (Fig. 2d, e). Subsequently, tethering GFP-RNaseH1 at the LacO site led to a significant reduction in R-loop signal^14^, indicating suppression of R-loops by artificial recruitment of an R-loop resolvase. Intriguingly, tethering GFP-SMARCAD1 resulted in a reduction of R-loop intensity comparable to that observed with GFP-RNaseH1 (Fig. 2d, e), suggesting that SMARCAD1 can directly suppress R-loop accumulation.

### SMARCAD1 senses R-loops through its N-terminus and resolves them via its C-terminus ATPase domain, promoting fork symmetry

Based on our *in vitro* studies, the ATPase domains are responsible for R-loop binding. To test if SMARCAD1 regulates R-loop levels *in vivo* through its ATPase-dependent nucleosome remodeling activity, we used the previously characterized catalytically dead point-mutant K528R that can interact with replication forks but is defective in nucleosome remodeling activity^42,48^ (Extended Data Fig. 3d). Even though data mining had identified SMARCAD1 as a potential R-loop binding protein associated with active replication forks (Extended Data Fig. 1a, b), it is unclear if SMARCAD1 can sense R-loops in the vicinity of forks. To investigate this, we also used our previously characterized NΔ-SMARCAD1 mutant, which lacks amino acids 1-136 and, therefore, is unable to associate with the replication fork^42^. We further validated our previous observation of SMARCAD1 interaction with active forks, previously shown with PCNA^42^, by performing a proximity ligation assay (PLA) between SMARCAD1 and nascent DNA labeled with 5’-ethynyl-2’-deoxyuridine (EdU). We compared wild-type SMARCAD1 with two SMARCAD1 mutants: NΔ-SMARCAD1 and K528R- SMARCAD1 mutants (Fig. 3a). The PLA results showed a drastic reduction in signal for the NΔ-SMARCAD1 mutant, indicating a loss of interaction with replisomes (Fig. 3b). In contrast, the K528R-SMARCAD1 mutant exhibited no such reduction, similar to wild-type, confirming that, while the NΔ-SMARCAD1 mutant disrupts the interaction with replisomes, the K528R-SMARCAD1 mutant does not. To further validate these findings, we performed PLA in the presence of hydroxyurea (HU) to examine SMARCAD1 dissociation from stalled replication forks, as previously reported^42^ and supported by iPOND-SILAC-MS (Extended Fig. 1b and Table 1). Consistent with SMARCAD1 role at active replication forks^42^, we observed a significant reduction in PLA signal under all HU conditions (Fig. 3b), corroborating its dissociation from stalled forks.

**Figure 3.**
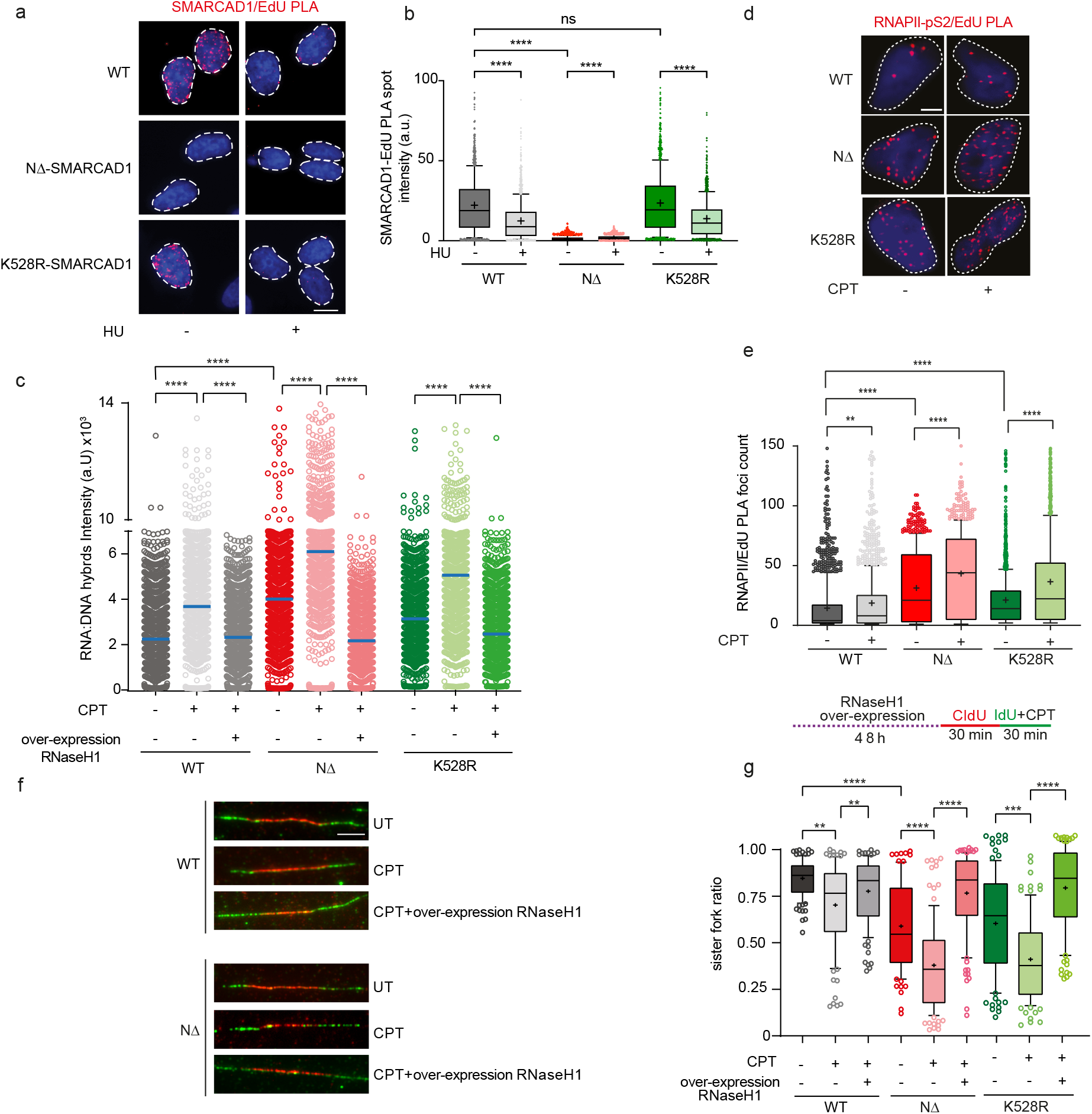
SMARCAD1 association via its N-terminus and binding R-loops via its ATPase in C-terminus, both are essential for R-loop resolution and maintaining fork symmetry. **a.** Representative images of PLA between SMARCAD1 and EdU in indicated cell lines and conditions. Scale bar, 10 μm. **b.** Boxplot representation of the quantification of high-content PLA spot count between SMARCAD1 and EdU of the indicated conditions. HU treatment was performed at 1 µM for 1 h. n(WT UT) = 1000, n(WT CPT) = 1000, n(NΔ UT) = 1000, n(NΔCPT) = 1000. ns = non significant, ****=p≤0.0001, one-way analysis of variance Kruskal-Wallis test, followed by Dunn’s correction for multiple testing. Source numerical data available in Source Data. **c.** Quantification of high-content DNA-RNA hybrids intensity of the indicated conditions. CPT treatment was given 1 h before the assay at 25nM. n(WT UT) = 1717, n(WT CPT) = 1700, n(WT CPT oeRNaseH1) = 1703, n(NΔ UT) = 1717, n(NΔ CPT) = 1699, n(NΔ CPT oeRNaseH1) = 1689, n(K528R UT) = 1000, n(K528R CPT) =1000, n(K528R CPT oeRNaseH1) = 1000 cells. ****=p≤0.0001, one-way analysis of variance Kruskal-Wallis test, followed by Dunn’s correction for multiple testing. Source numerical data available in Source Data. **d.** Representative images of PLA between RNAPII-pS2 and EdU in indicated cell lines and conditions. Scale bar, 10 μm. **e.** Boxplot representation of the quantification of high-content PLA foci count between RNAPII-pS2 and EdU of the indicated conditions. CPT treatment was given 1 h before the assay at 25nM followed by a short pulse of 20 minutes of EdU. n(WT UT) = 1535, n(WT CPT) = 1487, n(NΔ UT) = 823, n(NΔ CPT) = 835. ****=p≤0.0001, one-way analysis of variance Kruskal-Wallis test, followed by Dunn’s correction for multiple testing. Source numerical data available in Source Data. **f.** Representative examples of sister fork asymmetry observed by sequentially labeling nascent replicating DNA. Scale bar, 5 μm. **g.** Top: schematics of experimental set-up with Dox-inducible over-expression of RNaseH1, CPT treatment during IdU labeling at 25 nM and sequential labeling of nascent DNA. Bottom: boxplot representation of the quantification for sister fork ratio (ratio shorter/longer IdU tracks) of the indicated conditions from (**f**). n(WT UT) = 237, n(WT CPT) = 198, n(WT CPT oeRNaseH1) = 211, n(NΔ UT) = 186, n(NΔ CPT) = 202, n(NΔ CPT oeRNaseH1) = 210, n(K528R UT) = 200, n(K528R CPT) = 200, n(K528R CPT oeRNaseH1) = 200 replicative sister forks. ns = non significant, **=p≤0.01, ****=p≤0.0001, one-way analysis of variance Kruskal-Wallis test, followed by Dunn’s correction for multiple testing. Source numerical data available in Source Data.

Using high-content quantitative imaging-based cell cytometry analysis (QIBC) we confirmed significant R-loop accumulation in both the K528R and NΔ-SMARCAD1 mutants (Fig. 3c). This accumulation worsened with low doses of the DNA topoisomerase I inhibitor Camptothecin (CPT) which induces TRCs and R-loop accumulation^66–68^ (Fig. 3c). Particularly, we used a 1-hour treatment at 25nM of CPT, which enabled us to detect persistent R-loop accumulation, while avoiding detrimental induction of double-strand breaks^68^. To confirm specificity, we used a Dox-inducible over-expression construct for RNaseH1^25^ (Extended Data Fig. 4a) and we observed that the increased S9.6 signal in both SMARCAD1 mutants was restored to wild-type levels by RNaseH1 over-expression even upon CPT induction (Fig. 3c). This data suggests that both the ATPase activity and the association of SMARCAD1 to the replication machinery through its N-terminus are critical for its potential R-loop sensing functionality to facilitate their resolution.

To further test if SMARCAD1 mutants accumulate TRCs, we conducted a PLA between elongating RNA polymerase II (RNAPII-pS2) and nascent DNA labeled by EdU (Fig. 3d). We observed a significant increase in PLA foci in both the NΔ-SMARCAD1 and catalytically dead K528R-SMARCAD1 mutants compared to wild-type cells (Fig. 3e). Additionally, a significant enrichment of TRCs was observed in both mutants, with a higher increase in the NΔ-SMARCAD1 mutant than K528R in the presence of CPT treatment (Fig. 3e). These results support a role for SMARCAD1 in resolving R-loop-associated TRCs. The higher accumulation of TRCs in NΔ-SMARCAD1 suggests that its R-loop sensing functionality is critical in the vicinity of active forks to facilitate their resolution.

Given that R-loops impact the symmetric progression rate of bidirectional replication forks^69,70^, to further validate the hypothesis that SMARCAD1 functions as an R-loop sensor at active forks, we examined whether the impaired association of SMARCAD1 with the replisome (NΔ-SMARCAD1) or the absence of its ATPase activity (K528R-SMARCAD1) would affect the symmetric progression of bidirectional forks due to R-loop accumulation. We sequentially labeled nascent replicating DNA with 5’-chloro-2’-deoxyuridine (CldU in red) and 5’-iodo-2’-deoxyuridine (IdU in green), followed by measuring the ratio between the shorter and longer IdU tracks of each bidirectional replication fork (Fig. 3f and Extended Data Fig. 4b). We observed that both the NΔ-SMARCAD1 and K528R mutants led to significantly increased fork asymmetry, which was further exacerbated by CPT treatment. This fork asymmetry was completely rescued by RNaseH1 over-expression, confirming that R-loop accumulation was the major cause of fork asymmetry in both mutants. (Fig. 3g).

Altogether, these data suggest that while the ATPase region of SMARCAD1 is essential for R-loop binding and processing to promote unhindered fork progression, its association with the replisome is equally critical for R-loop-mediated TRCs sensing to facilitate R-loop resolution and ensure smooth fork progression (Fig. 1b, c, j-l and Fig. 3a-g).

### SMARCAD1 promotes R-loop resolution during replication

Since SMARCAD1 seems to be involved in resolving R-loops associated with TRCs, we investigated if R-loops accumulated preferentially in S-phase in SMARCAD1 mutant cells, as R-loop enrichment can be cell cycle-dependent and higher during replication^71–73^. To assess if, in our conditions (1h, 25nM), CPT treatment would impact EdU incorporation, we used whole cell imaging resolution through QIBC analysis, which showed no clear differences in EdU intensity (Extended Data Fig. 4c). Subsequently, we monitored the accumulation of R-loops in NΔ-SMARCAD1 cells under untreated and CPT conditions, both in S-phase and non-S-phase. We observed a clear accumulation of R-loops in replicating cells upon CPT treatment, in both wild-type and NΔ-SMARCAD1 cells (Fig.4a). However, the accumulation of R-loops was notably higher during S-phase in NΔ-SMARCAD1 cells, suggesting that SMARCAD1 is required to process R-loops preferentially in replicating cells.

Furthermore, since RNaseH1 is the enzyme that cleaves *bona fide* DNA-RNA hybrids, we assessed whether the loss of RNaseH1 in NΔ-SMARCAD1 mutant shows epistatic or additive effects. This resulted in a more pronounced accumulation of DNA-RNA hybrids compared to the loss of either factor alone, suggesting a synergistic effect between these two independent pathways. (Fig. 4b, c). The increased DNA-RNA hybrids observed in NΔ-SMARCAD1 cells could also be resolved by exogenous treatment with RNaseH1 enzyme^74^ (Extended Data Fig. 4d). We measured the levels of DNA-RNA hybrids probed by the S9.6 antibody via dot-blot in NΔ-SMARCAD1 compared to wild-type cells, revealing a remarkably higher enrichment of *bona fide* RNaseH1-sensitive DNA-RNA hybrids in NΔ-SMARCAD1 cells (Fig. 4d).

**Figure 4.**
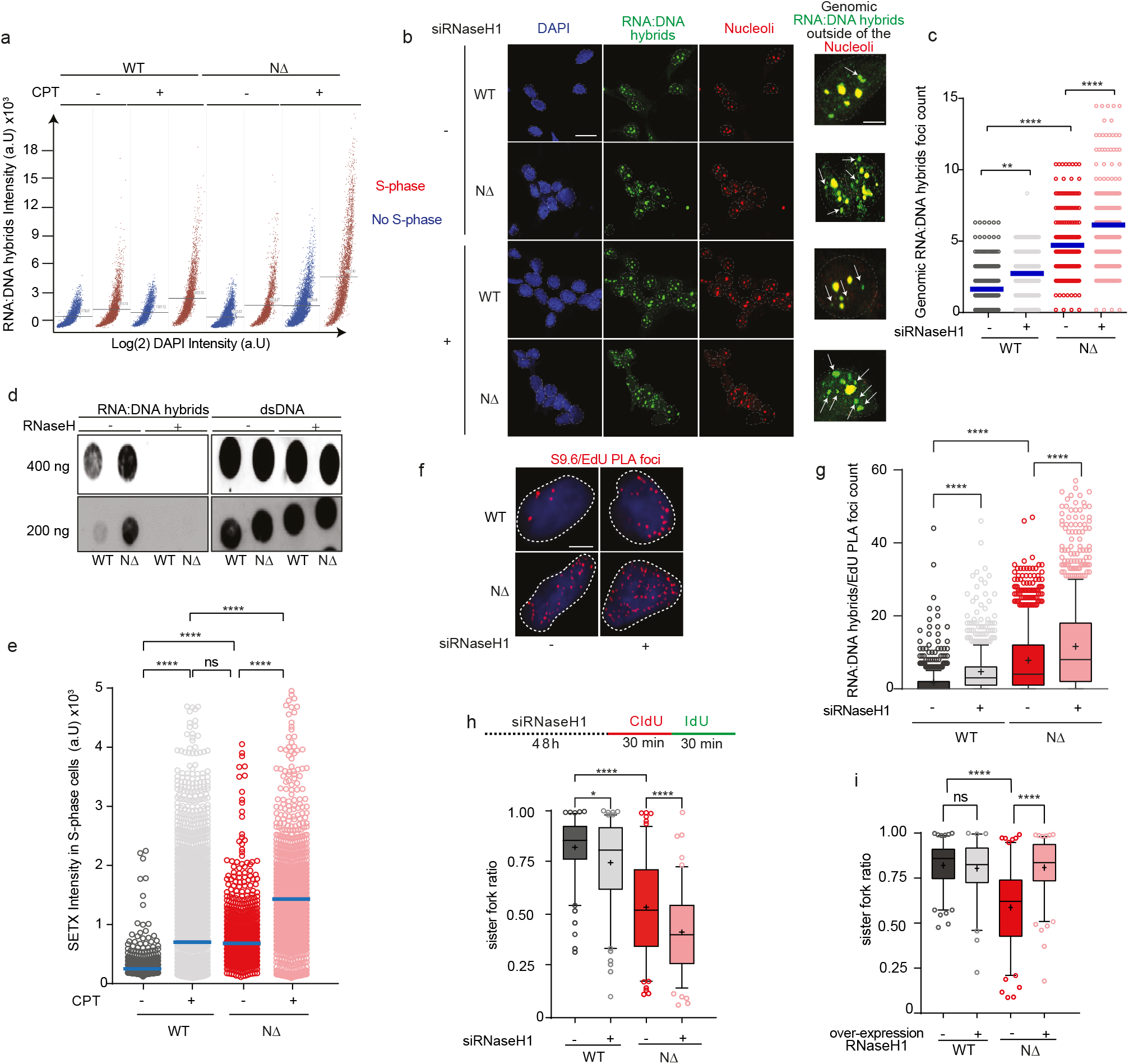
NΔ-SMARCAD1 accumulates R-loops during S-phase at active forks. **a.** QIBC plot of DNA-RNA hybrids intensity over DAPI intensity separating EdU+ cells and EdU-cells in indicated cell lines and conditions. CPT treatment was given 1 h before the assay at 25 nM. n(WT UT) = 1717, n(WT CPT) = 1700, n(WT CPT oeRNaseH1) = 1703, n(NΔ UT) = 1717, n(NΔ CPT) = 1699, n(NΔ CPT oeRNaseH1) = 1689 cells. **b.** Left: representative confocal images of R-loops immunofluorescent labeling using S9.6 antibody in MRC5 cells in indicated cell lines and conditions. RNaseH1 siRNA transfection was performed 48 h before the assay. Scale bar, 100 μm. Right: zoom highlighting DNA-RNA hybrids outside nucleoli (indicated by white arrows). Scale bar, 10 μm. **c.** Quantification for DNA-RNA hybrids foci count outside of the nucleoli of the indicated conditions using ImageJ. n(WT) = 243, n(WT siRNaseH1) = 240, n(NΔ) = 225, n(NΔ siRNaseH1) = 227 cells. ns = non significant, * = p<0.5, **=p≤0.01, ****=p≤0.0001, one-way analysis of variance Kruskal-Wallis test, followed by Dunn’s correction for multiple testing. Source numerical data available in Source Data. **d.** Dot-blot followed by western blot of indicated conditions in presence or absence of RNase H exogenous treatment. Uncropped membranes available in Source Data. **e.** Quantification of high-content immunofluorescence of SETX intensity in S-phase cells of the indicated conditions. CPT treatment was given 1 h before the assay at 25nM. n(WT UT) = 1518, n(WT CPT) = 3618, n(NΔ UT) = 1581, n(NΔ CPT) = 1749. ns = non significant, ****=p≤0.0001, one-way analysis of variance Kruskal-Wallis test, followed by Dunn’s correction for multiple testing. Source numerical data available in Source Data. **f.** Representative images of PLA between S9.6 and EdU in indicated cell lines and conditions. Scale bar, 10 μm. **g.** Boxplot representation of the quantification of high-content PLA foci count between S9.6 and EdU of the indicated conditions. n(WT UT) = 769, n(WT CPT) = 931, n(NΔ UT) = 769, n(NΔ CPT) = 991. ****=p≤0.0001, one-way analysis of variance Kruskal-Wallis test, followed by Dunn’s correction for multiple testing. Source numerical data available in Source Data. **h.** Top: schematics of experimental set-up with knock-down of RNaseH1 by siRNA and sequential labeling of nascent DNA. Bottom: boxplot representation of the quantification for sister fork ratio (ratio shorter/longer IdU tracks) of the indicated conditions. n(WT siCTRL) = 119, n(WT siRNaseH1) = 120, n(NΔ siCTRL) = 126, n(NΔ siRNaseH1) = 115 replicative sister forks. ns = non significant, ****=p≤0.0001, one-way analysis of variance Kruskal-Wallis test, followed by Dunn’s correction for multiple testing. Numerical data available in Source Data. **i.** Top: schematics of experimental set-up with Dox-inducible over-expression of RNaseH1 and sequential labeling of nascent DNA. Bottom: boxplot representation of the quantification for sister fork ratio (ratio shorter/longer IdU tracks) of the indicated conditions. n(WT) = 121, n(WT oeRNaseH1) = 122, n(NΔ) = 122, n(NΔ oeRNaseH1) = 122 replicative sister forks. ns = non significant, ****=p≤0.0001, one-way analysis of variance Kruskal-Wallis test, followed by Dunn’s correction for multiple testing. Numerical data available in Source Data.

We further tested if SETX, a primary resolvase of R-loops known to be specific for S-phase, as observed in human and budding yeast^17,72^, accumulates significantly in NΔ-SMARCAD1 to process enhanced R-loops. As expected, we observed an increase in SETX levels in wild-type S-phase cells upon CPT treatment, consistent with its functions (Fig. 4e). Interestingly, unchallenged NΔ-SMARCAD1 cells showed a significantly higher enrichment of SETX intensity, similar to the increased S9.6 accumulation or TRCs accumulation (Fig. 3a-c), suggesting an increase in R-loops. This SETX enrichment was further enhanced upon CPT treatment in NΔ-SMARCAD1 cells compared to wild-type CPT-treated cells (Fig. 4e). These findings imply that loss of SMARCAD1 function does not impact recruitment of SETX, and that SMARCAD1 might independently play a crucial role in processing R-loops, particularly during S-phase.

Given the preferential accumulation of R-loops during the S-phase of the cell cycle, we investigated whether this phenomenon occurs at the site of active replication forks. To visualize this link, we optimized a PLA between R-loops, using the S9.6 antibody, and nascent replicating DNA, labeled with a short pulse of EdU (Extended Data Fig. 4e). We conducted the S9.6/EdU PLA in the presence of RNaseH1 knock-down, aimed at increasing the levels of R-loops. Even in untreated conditions, we observed a significant increase in PLA foci in NΔ-SMARCAD1 cells, which was further elevated by the loss of RNaseH1 (Fig. 4f, g). Furthermore, we measured the fork asymmetry in NΔ-SMARCAD1 cells upon RNaseH1 knock-down and, according to our results, this led to an exacerbated asymmetric fork progression (Fig. 4h), while over-expression of RNaseH1 rescued this phenotype (Fig. 4i).

These results collectively suggest that conflicts accumulate in SMARCAD1 mutants in the proximity of replication forks, leading to asymmetric fork progression. Together, these data indicate that SMARCAD1 acts as a sensor of R-loops and potentially processes them independently in the proximity of active replication forks.

### SMARCAD1 processes R-loops independently of its role in PCNA homeostasis at replication forks

Previously, we showed that SMARCAD1 associates with active forks and promotes the stability of unperturbed as well as restarted stalled forks by restoring PCNA levels at forks^42^. We showed that loss of 53BP1 could rescue levels of PCNA in absence of SMARCAD1 which reduced the high turnover of PCNA by preventing its excessive unloading at active forks^42^. We wondered whether the observed processing of R-loops could be attributed to SMARCAD1 function in fine-tuning PCNA levels or rather it represents a unique function.

To dissect this function, we measured fork asymmetry in WT and NΔ-SMARCAD1 mutant cells upon knock-down of 53BP1 (Extended Data Fig. 5a). We observed that loss of 53BP1 could not rescue the fork asymmetry defects in NΔ-SMARCAD1 cells (Extended Data Fig. 5b). As previously shown^42^, NΔ-SMARCAD1 cells exhibit a marked fork progression defect, which can be rescued by 53BP1 knockdown. Indeed, our analysis confirmed that 53BP1 knock-down improves fork progression rates but does not correct the fork asymmetry phenotype (Extended Data Fig. 5b, c). This suggests that the R-loop accumulation causing fork asymmetry in SMARCAD1 mutants is independent of SMARCAD1 role in regulating PCNA levels at the replication fork.

To further confirm this, we tested SMARCAD1 mutants sensitivity towards CPT, which leads to an enhanced R-loop-associated TRCs accumulation, as well as fork asymmetry. Our clonogenic assays revealed significant sensitivity to low doses of CPT in SMARCAD1 mutants compared to wild-type cells (Extended Data Fig. 5d). Although 53BP1 knock-down in SMARCAD1 mutants rescues sensitivity to other replication-stress-inducing drugs, such as cisplatin and olaparib^42^, it did not rescue CPT sensitivity (Extended Data Fig. 5d). These results suggest that the observed phenotypes of R-loop-associated TRCs, fork asymmetry and CPT cellular sensitivity are not due to loss of PCNA homeostasis but could be linked to a unique role of SMARCAD1 in resolving conflicts at active replication forks.

### SMARCAD1 mutant cells accumulate R-loops preferentially at replication origin-proximal regions independently of replication directionality

To assess how SMARCAD1 regulates the processing of R-loops genome-wide, we examined the R-loop profile across the genome in NΔ-SMARCAD1 cells compared to wild-type. We confirmed the specificity of S9.6 immunoprecipitation for DNA-RNA hybrids, sensitive to RNaseH1, using DNA-RNA immunoprecipitation (DRIP)-sequencing at various genomic loci (Fig. 5a and Extended Data Fig. 6a). Genome-wide analysis revealed clear enrichments of R-loop signals in wild-type cells, which were significantly exacerbated in NΔ-SMARCAD1 cells (Fig. 5a). To confirm reproducibility, we plotted the DRIP peaks for each condition from two independent replicates. This showed a consistent and comparable signal across both replicates, prompting us to use average coverage for our further analysis (Extended Data Fig. 6b). Notably, the spread of RNaseH1-sensitive DRIP-seq signal was significantly more extensive in NΔ-SMARCAD1 cells compared to wild-type, encompassing 8305 genomic sites with 5-fold higher R-loop enrichment, compared to 1909 such regions in wild-type (Fig. 5b). These regions accumulate R-loops as defined by their specific immunoprecipitation with S9.6 that is not observed in the RNaseH1 treated samples in the DRIP-seq experiment. The peak calling was done using the RNase H1-treated sample as control to be certain that the analysis is done with confirmed DNA-RNA hybrids that are fully suppressed by treating with RNaseH1. The 8305 regions analyzed are enriched in R-loops in NΔ-SMARCAD1 cells 1.5-fold (p value <0.05) above the WT. These regions predominantly comprised intragenic regions, with over 60% of R-loops in NΔ-SMARCAD1 mutants augmented in genic regions, with 84% of these regions corresponding to protein-coding genes (Fig. 5c). We re-analyzed our RNA sequencing (RNA-seq)–based transcriptome analysis performed in these cell lines previously^42^ and cross-compared it with our DRIP-Seq analysis. We differentiated between the expression of genes with DRIP signal enrichment in NΔ-SMARCAD1 cells and the expression of other genes (the rest of protein coding genes with expression values >0). This analysis revealed a significant difference in expression between these two groups (Extended Data Fig. 6c), suggesting that in the NΔ-SMARCAD1 mutant, R-loops predominantly form at highly expressed protein-coding genes, similar to the WT condition.

**Figure 5.**
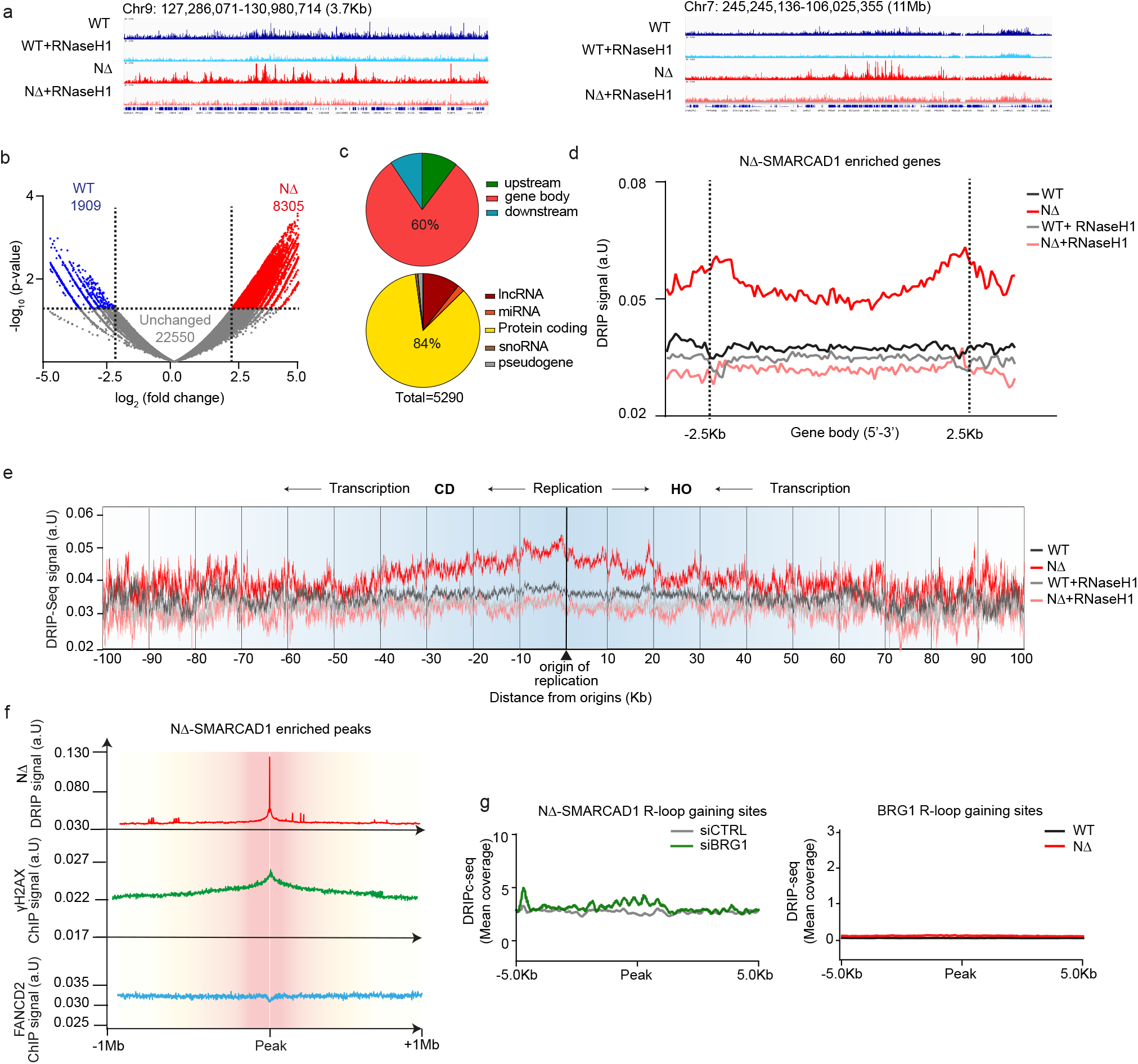
SMARCAD1 accumulates R-loops at highly-expressed coding gene regions and origin of replication. **a.** Representative screenshots of genomic regions showing DRIP-seq signal from WT (blue) and NΔ-SMARCAD1 (red) cells treated (light) or not (dark) with RNaseH1 *in vitro*. **b.** Volcano plot showing differential enrichment between WT (blue) and NΔ-SMARCAD1 (red). 1.5 FC and 0.05 p-value were set as threshold. **c.** Distribution of R-loop-enriched regions in NΔ-SMARCAD1 along genes (top) and in different gene classes (bottom). **d.** Distribution of DRIP-seq signal (average coverage) along protein-coding genes in WT (black) and NΔ-SMARCAD1 (red) samples treated (light) or not (dark) with RNaseH1 *in vitro*. **e.** DRIP-seq signal aligned with a OK-seq^93^ to determine the accumulation of R-loops around origins of replication. Windows of 10 Kb have been plotted using the going from –1 Kb to +9 Kb compared to the TSS of each window, up to 100 Kb. **f.** Distribution of NΔ-SMARCAD1 DRIP-seq signal (red) and the ChIP-seq signals ± 1 Mb γH2AX (green) and FANCD2 (blue) from wild type cells around the R-loop-enriched regions in NΔ-SMARCAD1. **g.** Right: Distribution of siCTRL (grey) and siBRG1 (green) DRIPc-seq signal in NΔ-SMARCAD1 R-loop-gaining sites. Left: Distribution of WT (black) and NΔ-SMARCAD1 (red) DRIP-seq signal in siBRG1 R-loop-gaining sites.

Furthermore, composite plots for gene bodies revealed substantial R-loop accumulation throughout the entire gene body in the NΔ-SMARCAD1 mutant compared to WT (Fig. 5d). To further characterize the genomic regions in which SMARCAD1 mutant accumulate R-loops, we cross-analyzed our DRIP-seq dataset with a short nascent strands sequencing (SNS-seq)^75^ that allows us to detect R-loop accumulation in the proximity of coordinates of replication origins. This analysis revealed a clear increase in RNaseH1-sensitive R-loop accumulation in SMARCAD1 mutant at origins of replication (Extended Data Fig. 6d). Moreover, we noted that 63% of the genes showing R-loop accumulation overlaps with origins (Extended Data Fig. 6e). Then, we wondered whether SMARCAD1 is required to resolve R-loops that accumulate in highly dynamic genomic regions near origins of replication, specifically in either head-on (HO) or co-directional (CD) orientations relative to actively progressing replication forks. To investigate this, we aligned our DRIP-seq dataset with a previous OK-seq^76^, and we plotted the DRIP-seq signal along protein-coding gene bodies located at distances from origins from 10-Kb to 100-Kb, to determine the average R-loop signal in the NΔ-SMARCAD1 mutant and WT, along with their respective RNaseH1-treated conditions. Notably, in the NΔ-SMARCAD1 mutant, R-loops accumulated primarily around the origins, regardless of HO or CD directionality (Fig. 5e). We further tested the possibility that enhanced R-loop accumulation near origins could alter origin firing in the NΔ-SMARCAD1 mutant, potentially causing fork progression defects. To do this, we used DNA combing technology to measure the frequency of origin firing. We observed that NΔ-SMARCAD1 did not show any significant alteration in origin firing frequency, as indicated by the length measured between origins and the number of origins fired similar to WT (Extended Data Fig. 6f). These results conclude that the defects observed in fork asymmetry in the NΔ-SMARCAD1 mutant are due to R-loop-mediated TRCs (Fig. 4h, i).

We previously provided evidence that FANCD2 functions together with the SWI/SNF chromatin remodeler BRG1 to promote R-loop resolution at stalled replication forks^22^. We wondered whether SMARCAD1 acted in the same process near blocked replication sites or if it is distinct. For this, we analyzed the correlation of R-loop-enriched sites in NΔ-SMARCAD1 with replication fork stalling detected by FANCD2 ChIP–seq^77^, and DNA damage revealed by γH2AX^78^, in wild-type cell lines as previously done for R-loop-enriched sites in BRG1-depleted cells^22^. Interestingly, R-loop accumulation in NΔ-SMARCAD1 cells correlated with DNA damage (γH2AX) but not with FANCD2-enriched sites (Fig. 5f), indicating that SMARCAD1 role in R-loop resolution differs from that described for SWI/SNF at blocked replication sites. Analysis of randomized DRIP-seq peaks further confirmed this lack of correlation (Extended Data Fig. 6g). This suggests that R-loops accumulate in NΔ-SMARCAD1 in regions highly prone to undergo DNA damage consistent with its role at actively progressing replication forks, particularly near origins. However, NΔ-SMARCAD1 R-loops do not accumulate at regions highly prone to undergo fork blockage in wild-type cells, given the lack of correlation between R-loop-enriched regions in NΔ-SMARCAD1 mutant and FANCD2-enriched regions in wild-type cells. This result suggests a different role of SMARCAD1 from that of SWI/SNF on R-loops, a conclusion further supported by cross-analysis of the DRIP-seq data from NΔ-SMARCAD1 MRC-5 cells with the DRIPc-seq data from siBRG1 K562 cells^22^. We found that loss of BRG1 did not lead to R-loop accumulation at the NΔ-SMARCAD1 R-loop-gaining regions and conversely, the NΔ-SMARCAD1 mutant did not accumulate R-loops in siBRG1 R-loop-gaining regions (Fig. 5g). Taken together, these findings suggest that SMARCAD1 and SWI/SNF function through distinct pathways at different stages of dynamic replication forks and in different genomic regions.

### R-loop-gaining regions specific to SMARCAD1 mutants correlate with hotspot mutations in germline cancer

Given the link between genome instability and cancer, and the relevance of R-loops as a source of genome instability, we next investigated whether the genome instability observed in SMARCAD1 mutants could be linked to R-loops and high mutation rates. In order to focus the analysis on the highly R-loop-enriched regions in SMARCAD1 mutants, we first re-analyzed the DRIP-seq coverage by generating metaplots of R-loop-gaining regions in NΔ-SMARCAD1 and wild-type cells. Notably, we found that the NΔ-SMARCAD1 mutant the R-loop peaks exhibited a >6-fold increase versus the WT, which showed relatively low coverage (Fig. 6a) consistent with a key role of SMARCAD1 in counteracting R-loop accumulation at regions in which they are rarely observed in wild-type cells. Next, we cross-referenced these R-loop-gaining sites in the NΔ-SMARCAD1 mutant with the Catalogue of Somatic Mutations in Cancer (COSMIC) database, assessing the abundance of single nucleotide variants (SNVs) at these sites (Fig. 6b). First, we computed SNV coverage across the genome and plotted the mean coverage along these sites, following previous methodologies ^22^. Remarkably, our analysis revealed that R-loop-enriched regions in the SMARCAD1 mutant exhibited increased SNV coverage in cancer (Fig. 6b). More specifically, we observed a strong enrichment of SNVs in single genes at R-loop-enriched sites in NΔ-SMARCAD1, while the correlation was weaker for insertions and deletions (Fig. 6c). These findings suggest that regions of *de novo* R-loop formation in the SMARCAD1 mutant are particularly susceptible to specific types of mutagenesis.

**Figure 6.**
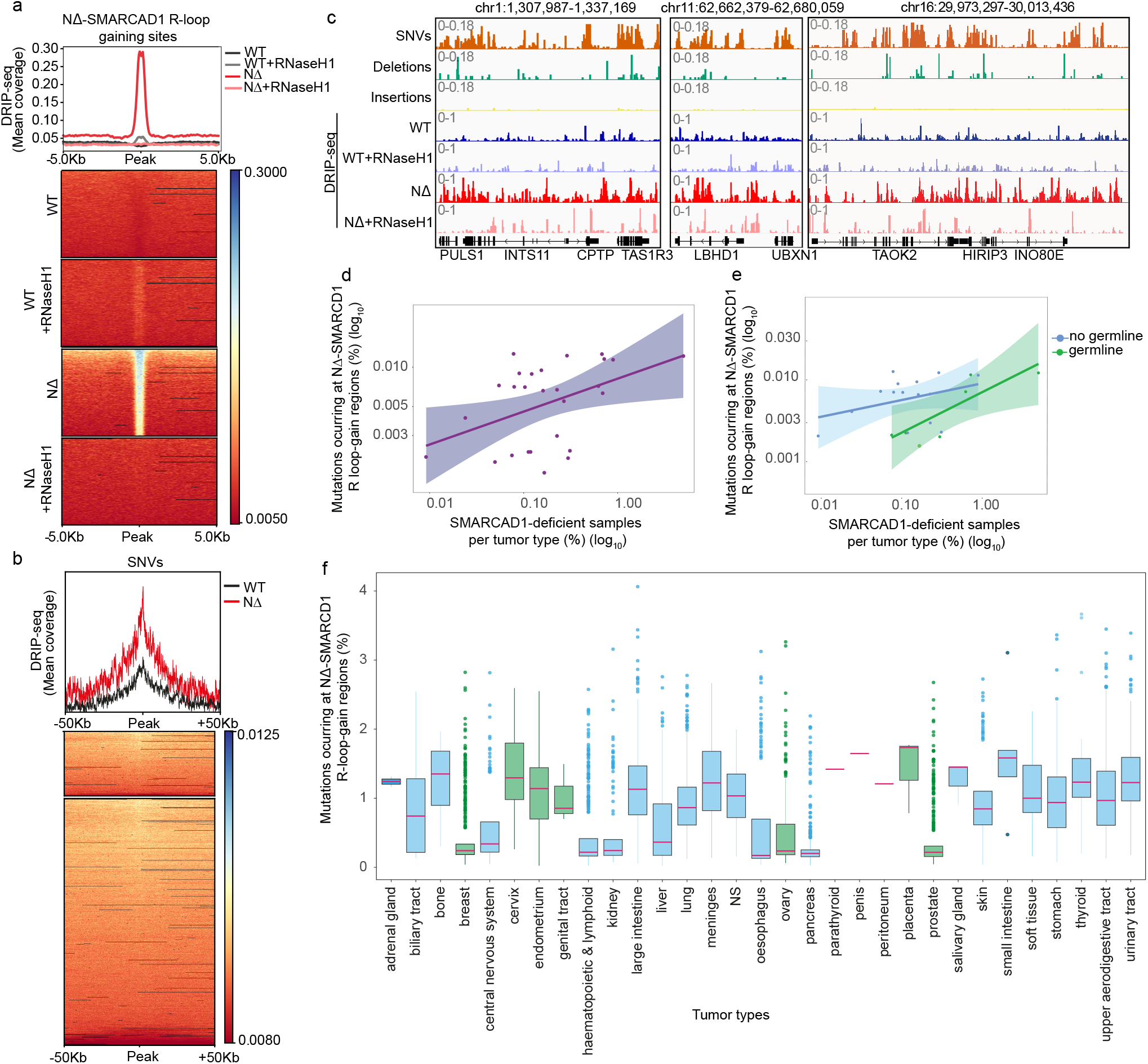
R-loop-gaining sites of SMARCAD1 mutant cells are mutation hotspots in tumors. **a.** Heatmap distribution of DRIP-seq signal of indicated conditions at NΔ-SMARCAD1 R-loop-accumulating regions **b.** Heatmap distribution of NΔ-SMARCAD1 R-loop accumulating regions and SNVs in WT and NΔ-SMARCAD1 **c.** Representative screenshots of genomic regions showing DRIP-seq signal from WT (blue) and NΔ-SMARCAD1 (red) cells treated (light) or not (dark) with RNH in vitro aligning it with SNVs signal (orange), deletions (green) and insertions (yellow). **d.** Correlation of SMARCAD1-deficinet tumors with NΔ-SMARCAD1 R-loop-gaining sites. **e.** Correlation between differential categories (non-germline in blue, germline in green) of SMARCAD1-deficient tumors with NΔ-SMARCAD1 R-loop-accumulating regions **f.** Analysis of mutation occurrence in different types of tumors in NΔ-SMARCAD1 R-loop-accumulating regions.

Further, considering the frequent mutation of chromatin factors, including SMARCAD1, in various cancers, it is plausible that mutagenesis of SMARCAD1 could contribute to extensive genome instability associated with tumorigenesis^79–84^. We assessed the frequency of mutations in specific sites per tumor sample and type. When correlating mutations in NΔ-SMARCAD1 R-loop-gaining regions with different tumor types exhibiting SMARCAD1 deficiency (Table 2), we found a mild (R^2^ = 0.20) yet significant (p = 0.020) correlation (Fig. 6d). We then categorized cancer types into germline tumors and non-germline tumors. Non-germline tumors showed no significant correlation, whereas germline tumors exhibited a strong (R^2^ = 0.71) and significant (p = 0.028) correlation (Fig. 6e). To explore the link between cancer-associated mutagenesis and R-loop accumulation in NΔ-SMARCAD1 mutants, we cross-referenced these data with the frequency of SMARCAD1-disruptive alterations, obtained though the COSMIC database (Extended Data Fig. 7). Interestingly, mutation frequencies associated with these sites positively correlated with SMARCAD1 alteration frequencies per cancer type, particularly in the reproductive system malignancies (Fig. 6f). Specifically, we noted a high percentage of mutations targeting R-loop-gaining sites in tissues such as cervix, endometrium, placenta, bone, small and large intestine, or meninges (Fig. 6f). These observations suggest a potential connection between SMARCAD1-mediated dysregulation of R-loop metabolism, genome instability, and the acquisition of mutations that could drive tumorigenesis. Further elucidation of this relationship may provide valuable insights into the role of SMARCAD1 in cancer development and the underlying mutagenic mechanisms involved.

**Table 2.**

| **Tumor** | **Typology** |
| --- | --- |
| Adrenal gland | Non-germline |
| Biliary tract | Non-germline |
| Bone | Non-germline |
| Breast | Germline |
| Central Nervous System | Non-germline |
| Cervix | Germline |
| Endometrium | Germline |
| Genital tract | Germline |
| Haematopoietic and Lymphoid tissue | Non-germline |
| Kidney | Non-germline |
| Large intestine | Non-germline |
| Liver | Non-germline |
| Lung | Non-germline |
| NS | Non-germline |
| Oesophagus | Non-germline |
| Ovary | Germline |
| Pancreas | Non-germline |
| Parathyroid | Non-germline |
| Penis | Non-germline |
| Peritoneum | Non-germline |
| Placenta | Germline |
| Prostate | Germline |
| Salivary gland | Non-germline |
| Skin | Non-germline |
| Small intestine | Non-germline |
| Soft tissue | Non-germline |
| Stomach | Non-germline |
| Thyroid | Non-germline |
| Upper aerodigestive tract | Non-germline |
| Urinary tract | Non-germline |

## DISCUSSION

In the present study, we mined and compared datasets of proteins that interact with R-loops and those associated with replication forks. From this analysis, we identified SMARCAD1, a unique chromatin remodeler that functions independently as a single 120 KDa protein, unlike other members of this family that are multi-subunit complexes^40,82,84,85^. This autonomy might allow SMARCAD1 to adapt swiftly to changing cellular environments and to efficiently manage chromatin remodeling tasks while travelling with the replication fork. SMARCAD1 is a multi-functional chromatin remodeler previously described in various roles, including heterochromatin inheritance preservation^41^, double stand break repair^40,86^, transcription regulation^38,39^ and active fork stability through PCNA homeostasis maintenance^42^. Here, we describe an additional and independent role of this chromatin remodeler.

Our data elucidate a direct interaction of SMARCAD1 with DNA-RNA hybrids and R-loops structures *in vitro*. The strong binding affinity, coupled with the competitive effect SMARCAD1 against RNaseH1 cleavage, suggests that SMARCAD1 plays a role in R-loop metabolism by directly interacting with the hybrid component. Furthermore, our *in vitro* data indicate that the ATPase/helicase domain is responsible for this interaction, while *in vivo* studies show that both the ATPase activity and the replication association domains of SMARCAD1 are critical for R-loop sensing and resolution. Although R-loops are unlikely to be compatible with the nucleosome formation^87^, it is possible that the role of SMARCAD1 in sensing and resolving R-loops, while ATPase-dependent, is independent of its chromatin remodeling function. The ATPase domain of SMARCAD1 may be required to destabilize the DNA-RNA hybrid structure of R-loops or facilitate their resolution *in vivo* through other resolvases, a possibility not excluded by our study. Further *in vitro* investigation of this activity could clarify whether the catalytic domains of ATP-dependent remodelers like SMARCAD1 play a distinct role in R-loop resolution, separate from their nucleosome remodeling activity, providing valuable insights into these mechanisms.

Interestingly, artificial tethering of SMARCAD1 suppressed R-loops at the LacO site to the same extent as artificial tethering of RNaseH1. This finding, in conjunction with the *in vitro* protection assay performed with SMARCAD1 and RNaseH1, suggests that these two proteins function on the same substrate. While we predict these enzymes to have distinct activity, RNaseH1 cleaves the RNA moiety from the DNA-RNA hybrid, the mechanism by which a chromatin remodeler like SMARCAD1 promotes resolution of the hybrid is unclear. However, our study provides evidence, using a unique combination of *in vitro* biochemical assays, functional genetic mutants, high-resolution quantitative imaging, and genomics technologies, that SMARCAD1 potentially plays an independent role in sensing and resolving R-loop/TRCs structures.

Here, we demonstrate a new factor for R-loop sensing and processing specifically at active replication forks, which operates independently of known factors that primarily function at blocked replication sites, such as SETX^17^, SWI/SNF^22^ or Fanconi Anemia^88,89^, or of the cell cycle independent RNaseH1^14^. We observe a significantly higher accumulation of R-loops in both SMARCAD1 mutants (NΔ-SMARCAD1 and K528R-SMARCAD1), underscoring the necessity of these domains for the *in vivo* sensing and resolution of TRCs and R-loops. Notably, this R-loop accumulation occurs preferentially during S-phase near active replication forks, leading to increased TRCs and asymmetric fork progression. These phenotypes are exacerbated by R-loop-inducing treatments, such as mild CPT or RNaseH1 knock-down, and are completely rescued by RNaseH1 over-expression, confirming the R-loops as the cause of the replication phenotypes.

Although the ATPase-containing C-terminal region of SMARCAD1 has been identified to directly interact with R-loops *in vitro*, the activity of sensing and resolving R-loops is insufficient without the involvement of its N-terminal domain, which links SMARCAD1 to the active replisome *in vivo*. This is evident from the observed phenotypes, which are either comparable to or slightly more severe in the NΔ-SMARCAD1 mutant compared to the point-mutant K528R-SMARCAD1. The latter still maintains its replisome association, unlike NΔ-SMARCAD1. Notably, critical phosphorylation sites have been identified in the N-terminus of SMARCAD1^28^ and its budding yeast homolog, Fun30, undergoes N-terminal phosphorylation in a cell cycle-dependent manner^90^. The NΔ-SMARCAD1 mutant lacks the initial 137 amino acids and many of the phosphorylation sites along with replication association. Together, this suggests that either post-translational modifications (PTMs) or replisome association through the N-terminus is critical for the proper functioning of SMARCAD1 in R-loop sensing. The ATPase function of SMARCAD1, which is essential for processing R-loops (this study) and maintaining active fork stability, is crucial for genome stability^42^.

Our genome-wide analysis revealed a substantial accumulation of R-loops in the NΔ-SMARCAD1 mutant compared to its wild-type, suggesting that SMARCAD1 plays a critical role in high-turnover processing of R-loops to prevent their buildup. This highlights SMARCAD1 unique function in facilitating R-loop processing, which is crucial for preventing fork stalling or enabling restart of blocked forks. Recent studies from our groups have shown that check-point activation at stalled replication forks leads to transient chromatin compaction and heterochromatinization, contrasting with the dynamics at active replication forks^91,92^. This might suggest that while DDR pathways are activated at blocked replication sites, the rate of R-loop resolution may be less critical compared to actively progressing forks, as evidenced by the enhanced accumulation of R-loops in SMARCAD1 mutant. Interestingly, R-loops in SMARCAD1 mutant cells are enriched at regions showing either head-on (HO) and co-directional (CD) conflicts in wild-type cells. This contrast with other cases like the human BRG1^22^ or yeast Sen1 whose mutations lead to R-loop accumulation preferentially at HO collision regions, known to be more detrimental to genome stability^10,12^. Recent studies also emphasize the importance of CD conflicts, which may necessitate fork restart^3^. The accumulation of both types of conflicts in SMARCAD1 mutant suggests that SMARCAD1 plays a more universal role in resolving R-loops near progressing replication forks. Taken altogether, we propose that SMARCAD1 acts early at active replication forks to facilitate the fast turnover of R-loops, which would otherwise persist and promote replication fork blocks. These blocks could become targets for specialized factors such as SETX^17^, SWI/SNF^22^ or the Fanconi Anemia pathway^88,89^ which may not always resolve them properly. The improper resolution of R-loops by these pathways is associated with the high mutation rates observed in germline tumors, leading to genome instability (Fig. 7).

**Figure 7.**
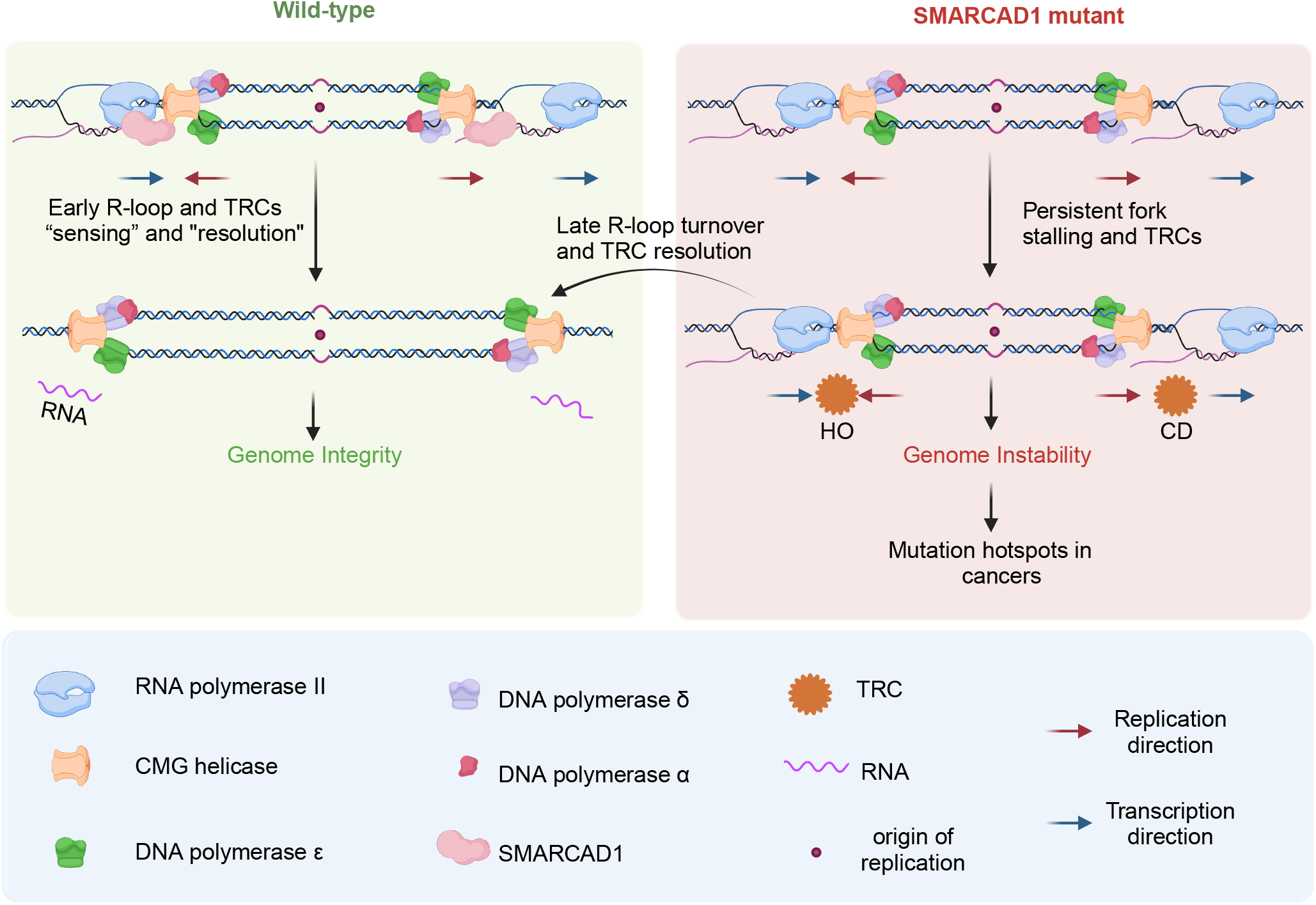
Dynamics of R-loop Resolution by SMARCAD1 at Actively Progressing Forks Working model. In wild-type and unchallenged conditions, SMARCAD1 travels with active replication forks, sensing R-loops associated with transcription-replication conflicts (TRCs) ahead of the forks via its flexible N-terminus domain. Upon detecting a conflict, the ATPase domain at the C-terminus directly interacts with the DNA-RNA hybrid of the R-loop, facilitating its rapid turnover to prevent fork stalling and activate the DDR. In SMARCAD1 mutants, conflicts are either not properly sensed or not quickly resolved, leading to enhanced R-loop accumulation at highly expressed and dynamic genomic regions, such as replication origins. These R-loops, which are turned over rapidly in wild-type and thus show minimal enrichment, appear *de novo* in SMARCAD1 mutants, resulting in increased genome instability that correlates with mutation hotspots in cancer. Figure was created with biorender.com.

Importantly, as SMARCAD1 is a frequently mutated chromatin remodeler in cancer, our study suggests a link between unresolved R-loop-mediated TRCs and cancer susceptibility. We observed that many R-loop-accumulating sites in the SMARCAD1 mutant are prone to DNA damage, with these sites showing a higher correlation with SNVs rather than larger insertions or deletions (indels). We have previously observed a similar correlation between R-loop-enriched sites in BRG1-depleted cells and mutation hotspots in cancer, including both SNVs and indels^28^. However, the R-loop accumulation prone regions in SMARCAD1 mutant conditions are distinct and it is likely that these regions may involve different error-prone repair pathways, such as translesion synthesis, which preferentially leads to SNVs rather than indels. This further supports that SMARCAD1 regulates R-loop differently to BRG1. Our data indeed suggest that SMARCAD1 prevents accumulation of dynamic R-loops observed at highly expressing genes, especially near origins of replication independent of their directionality in SMARCAD1-mutant. Interestingly, the R-loop regions accumulated in the SMARCAD1 mutant are frequently mutated and show a strong correlation with germline tumors (*i.e.* cervix, endometrium, ovarian, placenta). This is consistent with a role for SMARCAD1 in resolving R-loop-associated TRCs that may help explain the prevalence of SMARCAD1 mutations in human malignancies^93,94^ (Fig. 7). A deeper exploration of SMARCAD1’s multifaceted roles, particularly its autonomous and independent role in R-loop processing, could offer fundamental insights into the interplay between chromatin remodeling and R-loop biology. Additionally, this research could identify SMARCAD1 as a potential target for germline cancer treatment and new therapeutic strategies.

## MATERIALS AND METHODS

### Cell lines

All cell lines used are listed in Supplementary Table 1. MRC-5 human fibroblasts were cultured in a 1:1 ratio of Dulbecco’s modified Eagle’s medium (DMEM) and Ham’s F10 (Invitrogen) supplemented with 10% fetal calf serum (FCS, Biowest) and 1% Penicillin/Streptavidin (PS, Sigma-Aldrich) at 37 °C and 5% CO2 in a humidified incubator. U2OS 2-6-3 cell line was cultured in DMEM supplemented with 10% fetal calf serum (FCS, Biowest) and 1% Penicillin/Streptavidin (PS, Sigma-Aldrich) at 37 °C and 5% CO2 in a humidified incubator.

### Compounds and Materials

All compounds and drugs used are listed in Supplementary Table 2.

### siRNA transfection

siRNA smart pool for the indicated gene were purchased from Dharmacon and transfection were performed using lipofectamine RNAiMAX (Thermo Fisher) according to the manufacturer’s protocol for 2 consecutive days. Knock-down efficiency was checked by immunoblotting. All siRNA are listed in Supplementary Table 3.

### Over-expression plasmid for RNaseH1

For inducible over-expression of RNaseH1 we used plasmid pEBTet-BLAST-RNaseH1-myc/His, a kind gift from Dr. Manolis Papamichos-Chronakis from the University of Liverpool (Supplementary Table 4).

### DNA/RNA sequences

The list of nucleic acid sequences used in this study is included below. Sequences with a 6-FAM label are indicated with an asterisk at the corresponding position. Oligonucleotides (Supplementary Table 5) were purchased from IDT, resuspended, and their concentrations were determined by absorbance at 260 nm. DNA substrates for binding assays were prepared by annealing the correct oligos in a 1:1(:1) ratio in the following combinations: oligos 2 and 3 for dsDNAshort; oligo 1 and 2 for DNA-RNA hybrid; oligos 1, 4 and 5 for R-loopshort, ssRNA refers to unannealed oligo 1. Annealing was done in a PCR thermocycler set to heat to 90°C for 5 min and cool at 1°C min−1 to 4°C. Annealed substrates were verified on a 6% DNA retardation gel (Thermo Fisher) and stored at -20°C. The dsDNAlong is a 621 bp double-stranded DNA (used for AFM) that was expressed in *E. coli* and purified as previously described^51^.

### Protein expression and purification

All in vitro assays used dephosphorylated wild-type human SMARCAD1 isoform 2 with a naturally occurring V974A polymorphism (referred to as SMARCAD1 throughout the manuscript). SMARCAD1 protein was expressed and purified as previously described^34^. In brief, a pACEBAC1 plasmid containing the SMARCAD1 sequence with N-terminal 6-His tag was expressed in Sf9 cells. Cells were harvested and lysed via sonication. SMARCAD1 protein was purified from cell lysate using column chromatography. After nickel and cation exchange columns, the protein was mixed with Quick Calf Intestinal Alkaline Phosphatase (New England BioLabs) for 1 hour at room temperature to be dephosphorylated. The protein was then purified by nickel column followed by gel filtration over a S200 column. Purified protein was monitored throughout the purification process via SDS-PAGE. After column chromatography, pure fractions were pooled, and final concentration was determined via absorbance at 280 nm. Concentrated SMARCAD1 protein was stored at -80°C in a storage buffer of 100 mM KCl, 20 mM Hepes (pH 7.5), 2 mM TCEP, 1 mM AEBSF, and 10% glycerol.

### Electrophoretic mobility shift assay

Samples were prepared by mixing 50 nM DNA substrate with 1 µM SMARCAD1 in a buffer containing 50 mM Hepes (pH 7.5), 25 mM NaCl, 0.2 mg/ml BSA, 1 mM DTT, and 5% v/v sucrose for gel loading. Samples were incubated for 30 minutes at room temperature before loading onto a 5% native 59:1 polyacrylamide gel. Native gels were run with an electrophoresis buffer of 0.2x TBE at 4°C for 1 hour at a constant voltage of 150 V, and subsequently visualized using a GE Typhoon imager in fluorescence mode. This allowed visualization of the DNA substrates via the 5’-fluorescein label (FAM).

### Fluorescence polarization

Fluorescence polarization was monitored using a 5’-fluorescein label (FAM) on each of the substrates. DNA substrates at a final concentration of 10 nM were mixed with SMARCAD1 (0 to 1000 nM) in a binding buffer of 20 mM Tris (pH 7.5), 2 mM DTT, 1 mM EDTA, 0.01% CHAPS, and 0.01% NP-40. Samples were incubated at room temperature for 30 minutes, followed by measurement of fluorescence polarization as a function of SMARCAD1 concentration in a BMG Labtech CLARIOstar microplate reader. Data were analyzed in GraphPad Prism and fit to equation;

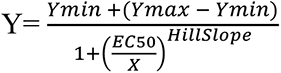

where Ymin and Ymax are the minimum and maximum signals, respectively, EC50 is the experimental binding constant, and X is the SMARCAD1 concentration. Each data point represents an average of three independent experiments ± the standard error.

### Protection Assay

Samples were prepared by mixing annealed R-loopshort with excess SMARCAD1 (final concentration 50 nM and 2 uM, respectively) in a buffer containing 50 mM Hepes (pH 7.5), 25 mM NaCl, 0.2 mg/ml BSA, 1 mM DTT. Samples were incubated at room temperature for 15 minutes and then complex formation was confirmed via fluorescence polarization. RNase H (purchased from NEB) was serial diluted in 1X RNase H reaction buffer (50 mM Tris-HCl, 75 mM KCl, 3mM MgCl2, 10 mM DTT) using an Opentrons OT-2 liquid handling robot, before being added to the samples and incubated at 20C for 20 minutes. Reactions were quenched with 5 mM EDTA, 0.5 mg/ml proteinase K, and 3.2% glycerol (for gel loading), before being incubated at 37C for 10 minutes. Samples were run on a 6% 29:1 native gel, with an electrophoresis buffer of 0.2x TBE for 40 minutes at a constant voltage of 100 V, and subsequently visualized using a GE Typhoon imager in fluorescence mode. This allowed visualization of the 5’-fluorescein label (FAM) on the RNA oligo. Analysis of the imaged gel was completed by quantifying the bands using ImageQuant software and graphed in Prism. Gels were also stained with Ethidium Bromide to visualize DNA.

### Preparation of R-looplong substrate

The R-looplong substrate was generously provided by the Fouquerel Lab (UPMC Hillman Cancer Center). In vitro transcription (IVT) for R-loop formation with the pFC53-mAIRN plasmid, as well as subsequent linearization and purification of the R-looplong substrate was completed as previously described^51^.

### Atomic Force Microscopy

For AFM with the R-loopshort, samples were imaged in buffer. AFM slides were prepared by freshly cleaving the mica, treating with APTES for 30 minutes, and rinsing with water. Slides were dried with filtered nitrogen gas. R-loopshort was diluted in 0M buffer (20 mM Tris-HCl pH 7.5, 1 mM EDTA) to a final concentration of 10 nM. Immediately after dilution, sample was applied to the APTES mica slide, rinsed with 0M buffer, followed by imaging in 0M buffer. Images were collected in AC mode on a NanoWizard 3.0 (Bruker) using a USC F0.3, k0.3 cantilever. All other AFM samples were imaged in air. LEtriDNA (90 nM) or R-looplong (65 nM) were incubated with SMARCAD1 (100 or 200 nM, respectively) in 0M buffer at room temperature for a minimum of 30 minutes. The reaction mixtures were diluted 50-fold in Imaging Buffer (20 mM HEPES, pH 7.5, 100 mM NaCl, 10 mM Mg(OAc)2) and immediately deposited onto a freshly cleaved mica surface. The samples were rinsed with water and dried with filtered nitrogen gas, followed by imaging in air. Images were collected in AC mode on a JPK/Bruker NanoWizard 3.0 with TAP300-Gold (Ted Pella) cantilevers. All images were captured at a resolution of 512 × 512 pixels at a scan rate of 1-3 Hz. AFM data was leveled, processed, and analyzed using Gwyddion software^95^. Particles were manually identified followed by extraction of the maximum height for each particle using Gwyddion software. Analysis and graphing were completed in GraphPad Prism (GraphPad Software, Boston, Massachusetts USA, www.graphpad.com). (Supplementary Table 7).

### Immunoblot

After lysis with RIPA buffer supplemented with protease inhibitor (Roche), samples were mixed with 2x Laemmli sample buffer (Supelco) and heated at 95 °C for 5 minutes. Samples were loaded on 4-12% NuPAGE Bis-Tris Gel (Novex life technologies) and transferred to a polyvinylidene difluoride (PVDF) membrane (0.45µm, Immobilon). Membranes were blocked with 5% BSA in PBS for 1 hour at room temperature and incubated with primary antibodies diluted in blocking buffer overnight at 4 °C, listed in Supplementary Table 6. Membranes were washed in 0.1% Tween-20 in PBS on the following day and subsequentially incubated with proper secondary antibodies coupled to near-IR dyes CF^TM^680/CF^TM^770. Membranes were visualized using and Odyssey CLx infrared scanner (LiCor).

### S9.6 antibody purification

S9.6 antibody used in immunostaining and dot-blot experiments was purified by GC lab as previously published^96^. Briefly, 800 ml of HB-8730 cellular supernatant was loaded in two columns respectively filled with 2 ml of Sepharose Protein A binding column, (GE Healthcare Chicago, IL, USA) and 2 ml of Sepharose Cl-4B protecting column, (GE Healthcare, Chicago, IL, USA), assembled one over the other. The columns were washed with phosphate buffer pH 8.0 and then the antibody was released from column with C-buffer pH 3.7 (68.8 mM citric acid, 35.9 mM Tris-Na-citrate), collecting the drops into Eppendorf tubes filled with 250 μl of Tris 1 M pH 8.5. After concentration and buffer exchange of positive fractions, the S9.6 antibody was quantified by Lowry Assay and checked by titration in immunofluorescence experiments to obtain the optimal concentration to be used. The S9.6 antibody used in DRIP-seq experiments was purified by AA lab from the HB-8730 hybridoma.

### DNA fiber analysis

Cells were sequentially labeled with 30µM CldU (MP Biochemicals) and 250µM IdU (Sigma-Aldrich) according to the schematic in each figure. After labeling, cells were collected and resuspended in PBS at 2.5 × 10^5^ cells/ml. Spreading and labeling of DNA was performed as previously described^91^.

### DNA combing

Cells were sequentially pulse labelled with 30 μM CldU (MP Biomedicals) and 250 μM IdU (Sigma-Aldrich) for 20 min each. Cells were collected, washed twice in PBS and resuspended in PBS at a concentration of 1.6 × 106 cells ml^−1^. DNA was extracted after encapsulation of cells in low-melting-point agarose blocks at 70,000 cells per plug and combed on silanized coverslips as described^97^. Detection of IdU and CldU labels was performed as described in the DNA fiber analysis procedure. Total DNA was labelled for 1h with anti-ssDNA antibody (AB_10805144, DSHB, 1:50), followed by 1h incubation in the dark with anti-mouse Alexa Fluor 350 (1:50) (Invitrogen). DNA fibers were then visualized and imaged as described above (DNA fiber analysis).

### Clonogenic survival assay

Cells were seeded in triplicate in 10cm culturing dish and treated with different concentrations of CPT for 4 hours and the washed and replaced with new medium. After 7 days colonies were fixed and stained in a mixture of 43% water, 50% methanol, 7% acetic acid and 0.1% Brillant Blue R (Sigma-Aldrich) and then counted. The survival was plotted as the mean percentage of colonies detected following the treatment and then normalized to the mean of the number of colonies in untreated samples.

### High-content immunostaining to detect DNA-RNA hybrids

For indirect immunofluorescence with S9.6 antibody against DNA-RNA hybrids in NT lab, cells were grown on coverslips at ∼70% confluency were either treated with 25nM CPT for 1 hour, or RNaseH1 was over-expressed by Dox-inducible plasmid transfection for 2 consecutive days, adding Dox 24 hours prior to the experiment. Cells were processed as previously described^85^. Briefly, cells were fixed in ice-cold methanol for 10 minutes at -20 °C followed by 1-minute permeabilization with ice-cold acetone. Cells were then washed three times in 4x SSC buffer and were blocked in 3% BSA for 1 hour in dark. Then, cells were incubated with S9.6 antibody at 1:250 dilution. Appropriate secondary antibody was used and DNA was stained using 2µg/µl of DAPI. Coverslips were then mounted using ProLong^TM^ Gold antifade mountant (Invitrogen). Slides were kept in dark at 4°C until imaged.

### LacO/LacR artificial tethering R-loop detection

For indirect immunofluorescence with S9.6 antibody in MSL lab, U2OS 2-6-3 cells were grown on coverslip and then transfected with fusion protein αGFP-LacR-mCherry^56^ and either with no additional plasmid, or with GFP-expressing plasmid, GFP-RNaseH1 plasmid and GFP-SMARCAD1 plasmid (Supplementary Table 4). Cells were then fixed as described above. Cells were incubated with S9.6 antibody a1:1000 dilution to allow only visualization of S9.6 signal at LacO array. Slides were treated as described above and kept in dark at 4°C until imaged. Images of fixed samples were acquired on a Zeiss AxioImager M2 widefield fluorescence microscope equipped with 63x PLAN APO (1.4 NA) oil-immersion objectives (Zeiss) and an HXP 120 metal-halide lamp used for excitation. Fluorescent probes were detected using the following filters for DAPI (excitation filter: 350/50 nm, dichroic mirror: 400 nm, emission filter: 460/50 nm), Alexa 488 (excitation filter: 470/40 nm, dichroic mirror: 495 nm, emission filter: 525/50 nm), or Alexa 647 (excitation filter: 640/30 nm, dichroic mirror: 660 nm, emission filter: 690/50 nm). Images were recorded using ZEN 2012 (blue edition, version 1.1.0.0) and analyzed in Image J (1.47v- 1.48v). Images were analyzed using ImageJ. LacO signal was masked and intensity of S9.6 was measured only within this region.

### High-content Proximity Ligation Assay (PLA)

PLA experiments were performed as previously described^91^. Briefly, cells were grown on coverslips pulse-labeled with EdU for 20 minutes. Cell were fixed in 4% Formaldehyde and permeabilized in 0.5% Triton-X in PBS for 10 minutes. For detection between RNAPII-pS2 and EdU, slides were then washed and blocked with 5% BSA in PBS for 1 hour at room temperature in dark. After two washes, freshly prepared Click-it reaction mix (2mM copper sulfate, 10µM biotin-azide and 100mM sodium ascorbate in PBS) was added to all samples at room temperature for 1 hour. Slides were incubated with primary antibodies at 4°C overnight. Following manufacturer’s protocol, slides were washed three times using buffer A (0.01M Tris, 0.15M NaCl and 0.05% Tween-20, pH 7,4) for 5 minutes each. Subsequentially, slides were incubated with Duolink in Situ PLA probes anti-mouse plus and anti-rabbit minus for 1 hour at 37°C in a humid chamber. After washes with buffer A, slides were incubated with Duolink ligation mix for 30 minutes at 37°C in a humid chamber and then incubated with Duolink amplification mix for exactly 120 minutes at 37°C. Slides were washed 3 times with buffer B (0.2M Tris and 0.1M NaCl in PBS) for 10 minutes each and one time with 0.01x buffer B. DNA was counterstained using DAPI for 15 minutes and slides were mounted with ProLong^TM^ Gold antifade mountant (Invitrogen). Slides were kept in dark at 4°C until imaged.

### Microscopic analysis of fixed cells and DNA fibers

High-content IFs, PLAs and DNA fibers were visualized with Metafer5 using 40X Plan-Neofluar 0.75NA air object. IFs were quantified using CellProfiler program, PLAs were quantified using MetaSystem and DNA fibers were quantified using ImageJ. All software used are listed in Supplementary Table 7.

### Dot-blot analysis of R-loops

Dot-blot was performed as described in^98^, with minor changes. Cells were seeded, collected and centrifuged at 300 x g for 5 minutes at 4°C. Cell pellets were incubated with cellular lysis buffer (10% NP-40, 2M KCl, 0.5M PIPES pH 8, nuclease-free water) and incubated on ice for 10 minutes. Nuclei were spun down at 500 x g for 5 minutes and resuspended in 400µl of ice-cold nuclear lysis buffer (10% SDS, 1M Tris-HCl pH 8, 0.5M EDTA, nuclease-free water). Nuclei were centrifuged again and incubated with 20mg/ml Proteinase K and incubated 3 hours at 55°C. Viscous DNA was sonicated with a 30s ON/30s OFF cycle for 10 minutes in a 4°C water bath. DNA was purified using elution buffer (1M Tris-HCl pH 8.5, nuclease-free water) and a mix of phenol:chloroform:isoamyl alcohol in a 25:24:1 ratio. After centrifugation at 12.000 x g for 5 minutes at 4°C, the aqueous phase was collected and extracted using 1 volume of chloroform. The new aqueous phase was then purified using 3M sodium acetate, glycogen and 70% ice-cold ethanol and spun down for 30 minutes at 4°C. Pellets were washed with isopropanol and the pellet was allowed to air dry for ∼30 minutes. Pellets were resuspended in water at 4°C overnight. Separately, a set of samples was treated either 5U of RNase H or 0.5U of RNase III or mock treated. Sampled were incubated at 37°C for 1 hour. A positively-charged nylon membrane was pre-washed using 2x SSC buffer. Then extracted R-loops, untreated and treated with RNase H and RNase III enzymes, were then washed with 6x SSC buffer. The membrane was dried for 5 minutes and the crosslinked using UV-A light at 1200µJ. Membrane was blocked using 5% milk in Tris with 0.05% Tween-20 (TBST) for 1 hour at room temperature. The membrane was then incubated with primary antibodies at 4°C overnight. The following day the membrane was washed three times with TBST and incubated with horseradish peroxidase (HRP)-conjugated secondary antibodies. ECL reagents were used to acquire the chemiluminescent signal.

### DRIP-seq

DNA-RNA hybrids were immunoprecipitated using the S9.6 antibody (hybridoma HB-8730), on purified genomic DNA enzymatically digested with HindIII, EcoRI, XbaI, SspI and BsrGI restriction As a control, samples were in vitro treated with RNaseH (New England BioLabs) as described^22,99^. Finally, eluted DNA was sonicated and size-checked on a 2100 Bioanalyzer (Agilent) and libraries constructed using the ThruPLEX DNA-Seq 6S kit (Rubicon Genomics) according to the manufacturer’s guidelines. Previous to the sequencing DRIP samples were tested for increased R-loop levels by DRIP-qPCR following standard procedures^100^

### DRIP-seq analysis

Sequenced paired-ends reads from DRIP-seq experiments were subjected to a quality control pipeline using the FASTQ Toolkit V.1.0.0 software (Illumina) and then mapped to the hg38 reference genome using the Rsubread V2.0.1 software package with unique=TRUE parameter^101^. Duplicated reads were marked using SAMtools V1.10^102^. Peak calling was performed with MACS2^103^, using RNH-treated sample as control. Regions covered by peaks in both replicates were retained and merged BEDtools^104^. For comparative purposes, WT and NΔ-SMARCAD1-resulting peaks were merged and combined when closer than 1kb using BEDtools^104^. The differential enrichment of these regions in each condition was performed using csaw V1.20.0 software package^105^. First we counted count reads in full genome using windowCounts() with bin = TRUE and width = 10000 parameters. Then normalization factors were calculated using normFactors(). Counts per peak were calculated using regionCounts for each condition and replicate and then normalized. After that, estimateDisp(), glmQLFit() and glmLRT() from edgeR package (v3.20.9), was used in order to calculate log2FC and p-value of the peaks. R-loop enriched regions were established selecting those peaks whose DRIP signal fold change was higher than 1.5 X and the (p-value) was lower than 0.05. Regions enriched in R-loops in NΔ-SMARCAD1 condition were annotated to genes retrieved from Ensembl release 86^106^ with ChIPpeakAnno V3.28.1 software package^107^. Promoter regions were established from the transcription start site (TSS) to 2 kb upstream and termination from transcription termination site (TTS) 2 kb downstream. Peaks were allowed to be annotated to more than one gene. OK-seq data and ChIP-seq data of SMARCAD1, γH2AX, and FANCD2 performed in K562 cells were retrieved from the European Nucleotide Archive (ENA) at EMBL-EBI under accession numbers PRJEB25180, PRJNA635402, PRJNA258172 and PRJNA473287 respectively. Files were mapped and processed similar to DRIP-seq data, being OK-seq data assigned to Watson and Crick strands using SAMtools V1.10^102^. Replicate fork directionality (RFD) track values were calculated as described previously for OK-seq data^76^. R-loop-enriched regions in NΔ-SMARCAD1 were stranded according to the genic annotation previously described, and then classified in CD or HO depending on its relative orientation with respect to the RF, given by the RFD. deepTools V3.4.3^108^ was used to calculate average coverages, generate CPM-normalized coverage profiles and metaplot images. Genome example regions were plotted using IGV V2.8.2 software^109^.

### SNVs and tumor metanalysis

Tumor mutation data was retrieved from the COSMIC database (http://cancer.ac.uk/cosmic) and data processed as indicated in ^28^. Mutation coverage plots were built using *deeptools*. Mutation frequencies at SMARCAD1-mutant R-loop gaining sites and SMARCAD1 alteration frequencies were computed and plotted using *R Studio* and *ggplot2* package.

### Statistics and reproducibility

Experimental data were plotted and analyzed using GraphPad Prism 9.4.1 (GraphPad Software). All box plots and violin plots show plain horizontal line representing the median and the “+” sign represents the mean of the dataset. The box contains the 25th to 75th percentiles of the dataset, the whiskers mark the 10th and 90th percentiles and values beyond these upper and lower bounds are considered outliers and marked with a dot. The number of samples analyzed per experiment are reported in the respective figure legends. All experiments were independently repeated at least two times with similar results obtained.

## Author contribution

NT and SU designed the study. SU: performed and analyzed all R-loops staining, QIBC assays, clonogenic survivals, PLAs, DNA fiber assays, dot-blot analysis, LacO-LacR artificial tethering experiments and generated figures. NMH: performed and analyzed *in vitro* biochemistry experiments including FP assays, protection assays, EMSA, AFM, generated the AlphaFold3 *in silico* prediction model, generated the corresponding figures and wrote section of their results in the manuscript. MES-O: analyzed DRIP-seq, generated data files, compared HO vs CD conflicts and DRIP-seq with different ChIP-seq data and generated the corresponding figures. AB-F: performed SNVs/indel analysis and tumor bioinformatic analysis and generated the corresponding figures. VG: performed and analyzed DNA fiber assay with si53BP1 and DNA combing assay. MLG-R: performed DRIP-seq experiments. CSYL: generated pipelines for high-content microscopy analysis and assisted SU in high-content imaging. CB: performed and analyzed SMARCAD1-EdU PLA. JM: purified S9.6 antibody used in immunostaining and dot-blot experiments. EMM: assisted SU with clonogenic survivals. GC: supervised JM. MSL: supervised SU with LacO-LacR tethering experiments and generated initial AlphaFold3 *in silico* predictions. KL: supervised NMH with *in vitro* experiments and data interpretation. AA: supervised MES-O and MLG-R with genome-wide data analysis and interpretations. NT: Supervised SU, VG, CSYL, CB and EMM; conceived and supervised the project. NT, SU, and AA wrote the manuscript. KL and NMH provided critical review and editing. All authors read, reviewed, and approved the final version of the manuscript.

## Acknowledgements

We thank Natalie Laspata and Elise Fouquerel (Fouquerel Lab, UPMC Hillman Cancer Center, Canada) for generously providing the purified and linearized R-looplong substrate for AFM analysis; Manolis Papamichos-Chronakis (Papamichos-Chronakis lab, Liverpool, UK) for kindly gifting plasmid pEBTet-BLAST-RNaseH1-myc/His for RNaseH1 over-expression; Haico van Attikum (van Attikum lab, LUMC, Netherlands) for pCMV6-AV-GFP- SMARCAD1 plasmid for LacO-LacR artificial tethering experiment, Daphne Boer (Luijsterburg lab, LUMC, Netherlands) for generating pLenti_CMV GFP puro and pLv_CMV- GFP-RNaseH1plasmids for artificial tethering experiment, and Sonia Barroso (Aguilera lab, CABIMER, Spain), for purifying and providing the S9.6 antibody used in DRIP-seq experiments. We thank the Genomic Unit of CABIMER (Seville, Spain) for DNA Sequencing. We thank current and former members of the Taneja lab.

---

**Extended Data Fig. 1.**
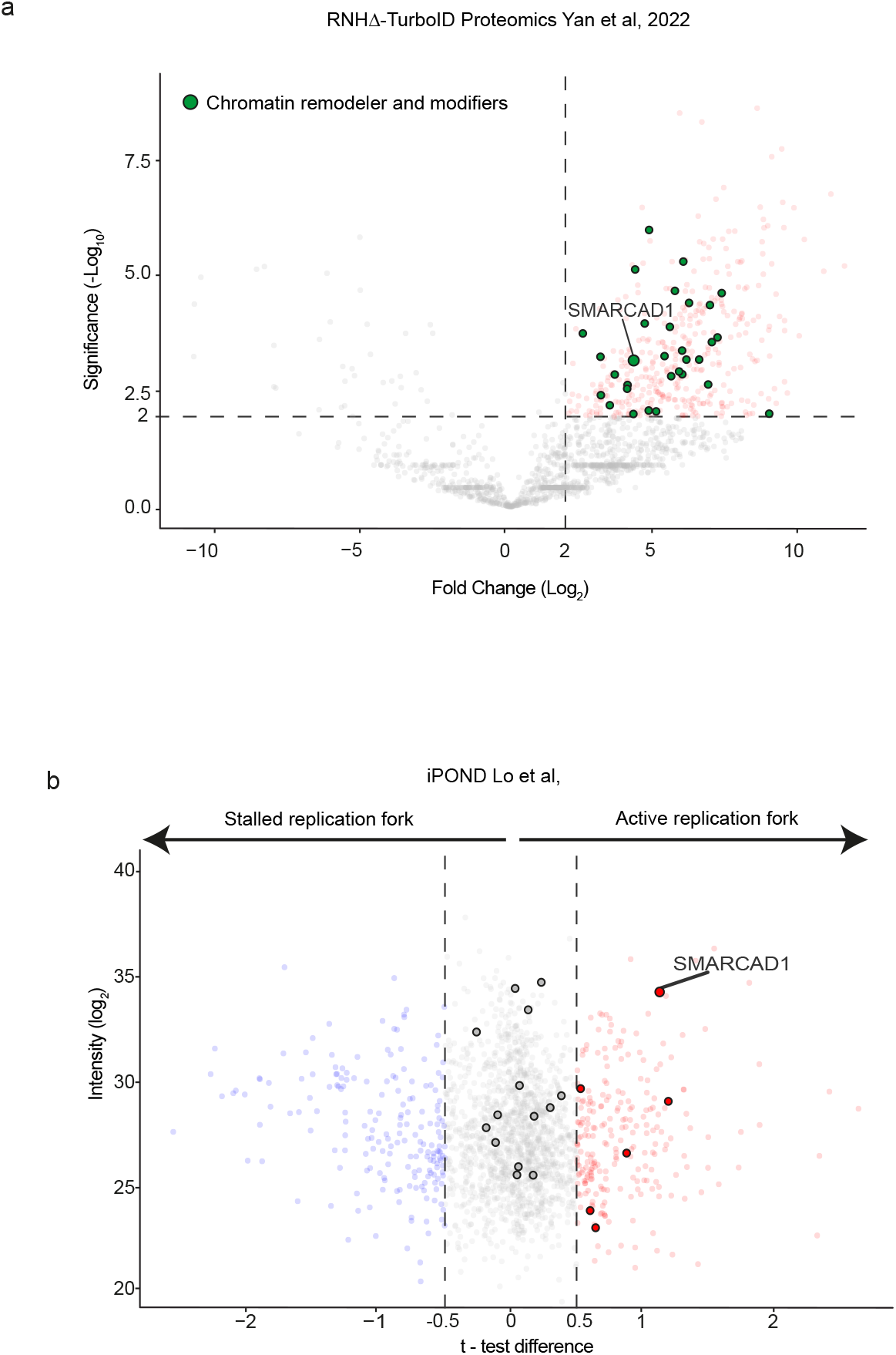
R-loop proximity and iPOND proteomics. **a.** Volcano plot of RNHΔ-TurboID proteomics from^46^ reanalyzed to highlight chromatin factors enriched. **b.** iPOND reanalysis from^42^ plotting chromatin factors enriched in RNHΔ-TurboID proteomics.

**Extended Data Fig. 2.**
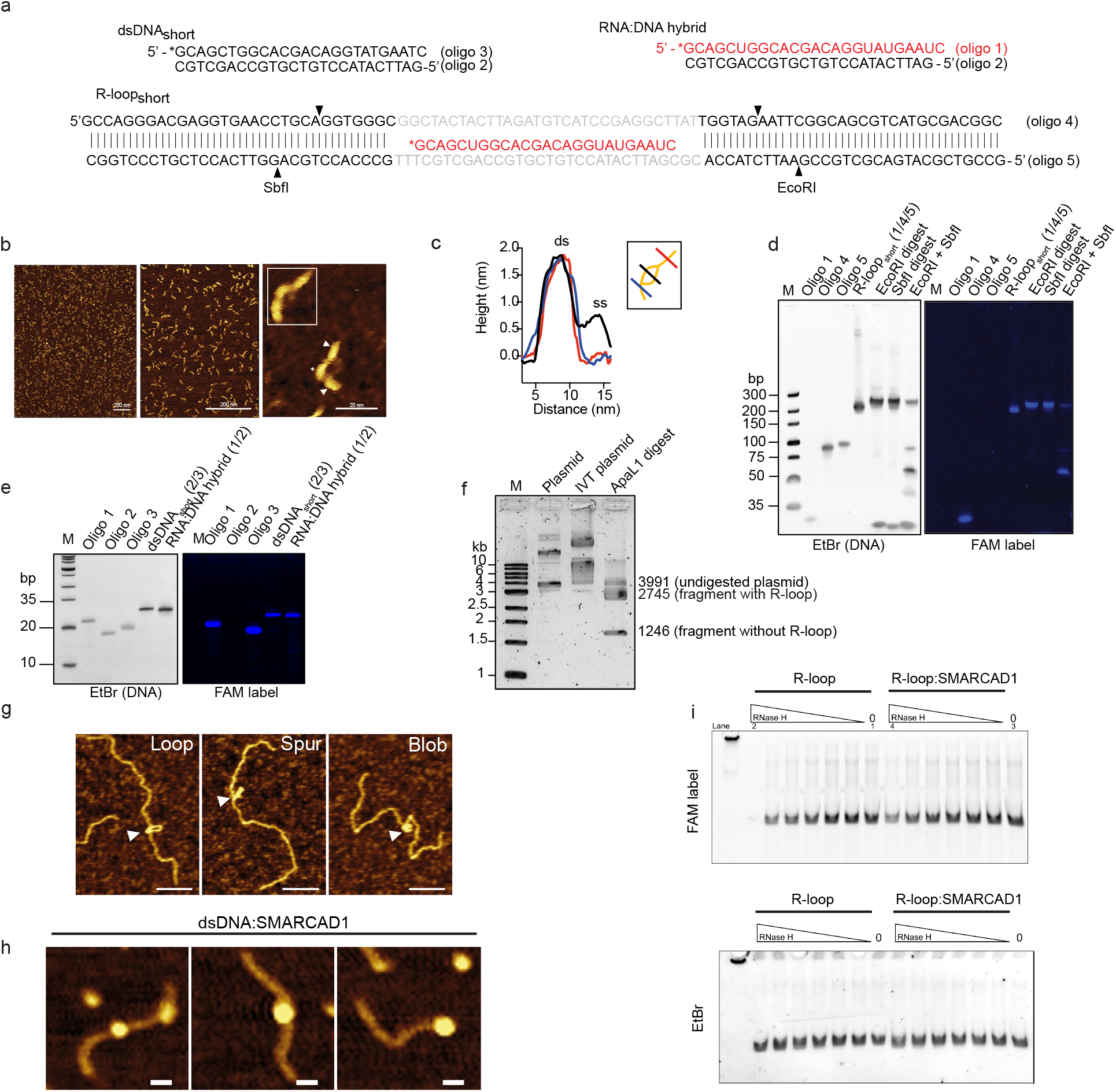
EMSA and AFM controls. **a.** Schematic and sequences of annealed substates. RNA bases are indicated in red text, FAM label is indicated as an asterisk at the corresponding position, and arrowheads indicate restriction sites. **b.** Representative AFM image of R-loopshort, arrowheads indicate dsDNA flanking the DNA-RNA hybrid (indicated by an asterisk). **c.** Height profile of the R-loopshort particle indicated with white square in **b**, along lines depicted in graphical inset. **d.** Validation of the R-loopshort substrate containing FAM labeled RNA (Oligo 1) via 6% 29:1 native gel, imaged with EtBr to visualize DNA and fluorescence to visualize the FAM label. Cleavage of the substrate by EcoRI and SbfI restriction enzymes confirms the presence of dsDNA flanking the R-loop. **e.** Validation of the DNA-RNA hybrid substrate containing FAM labeled RNA (Oligo 1) via 10% 19:1 native gel, imaged with EtBr to visualize DNA and fluorescence to visualize the FAM label. **f.** Agarose gel electrophoresis verifying R-loop formation in the pFC-mAIRN plasmid following IVT. Migration of un-transcribed plasmid (lane 1), transcribed plasmid (lane 2), and transcribed plasmid linearized by ApaL1 digest (lane 3) were analyzed. **g.** Representative AFM images of Rlooplong in various shapes (arrowheads). Scale bar, 100 nm. **h.** Representative AFM images of SMARCAD1 bound dsDNAlong. Scale bar, 20 nm. **i.** Uncropped gel images of the protection assay, imaged for fluorescence to visualize the FAM label (on RNA, Oligo 1) and EtBr to visualize DNA. Cropped lanes shown in Fig. 1h are numbered (1-4).

**Extended Data Fig. 3.**
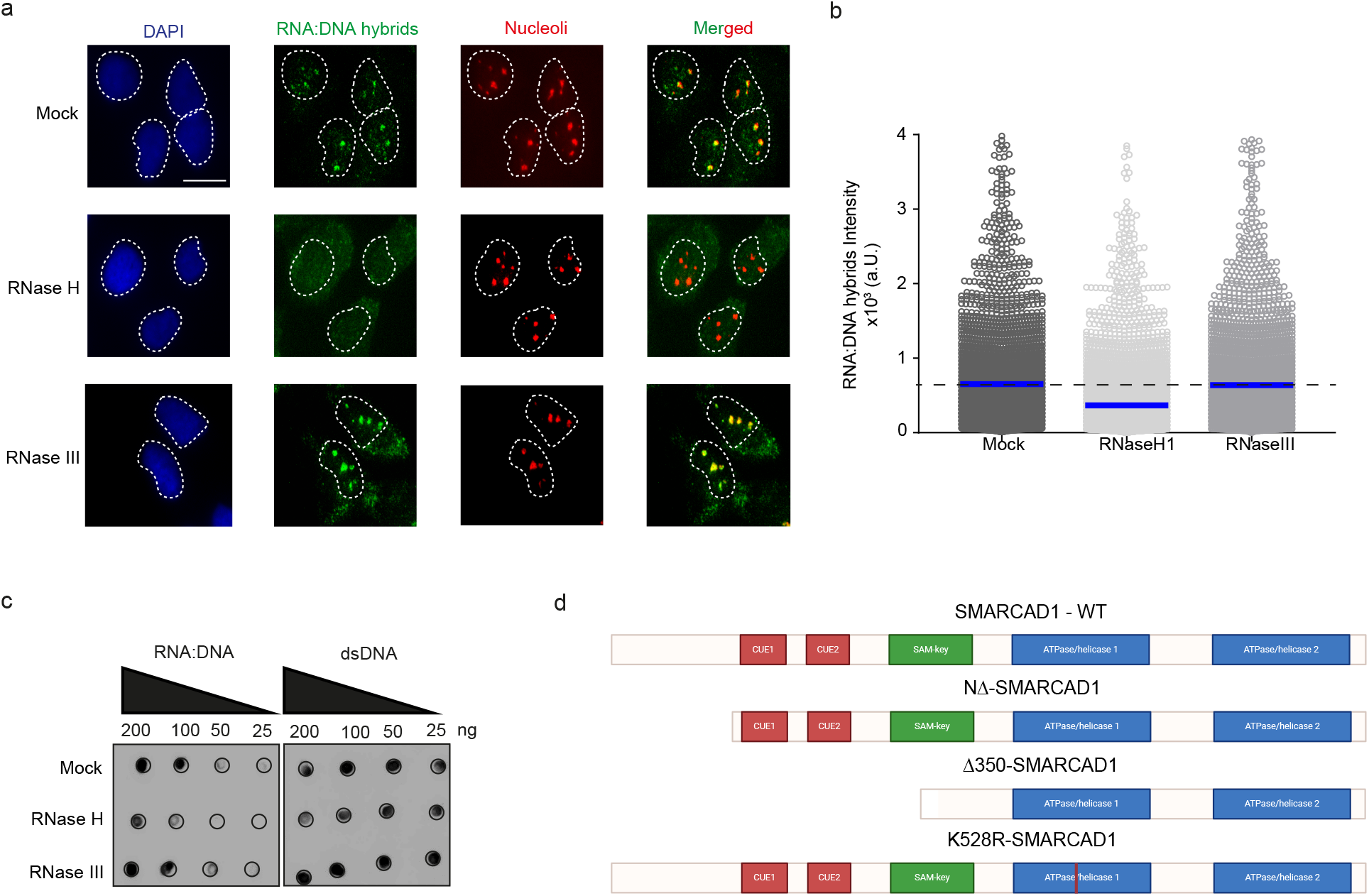
S9.6 antibody specificity for DNA-RNA hybrids. **a.** Representative images of DNA-RNA hybrids immunofluorescence of indicated conditions in MRC5 cells. Scale, 20µm. **b.** Quantification of high-content S9.6 intensity of the indicated conditions. n(mock) = 6251, n(RNaseH) = 8569, n(RNaseIII) = 5106. ns = non significant, ****=p≤0.0001, one-way analysis of variance Kruskal-Wallis test, followed by Dunn’s correction for multiple testing. Source numerical data available in Source Data. **c.** Dot-blot followed by western blot of indicated conditions. Uncropped membranes available in Source Data. **d.** Cartoon of SMARCAD1 mutants used for in vivo experiments. Figure was created with biorender.com.

**Extended Data Fig. 4.**
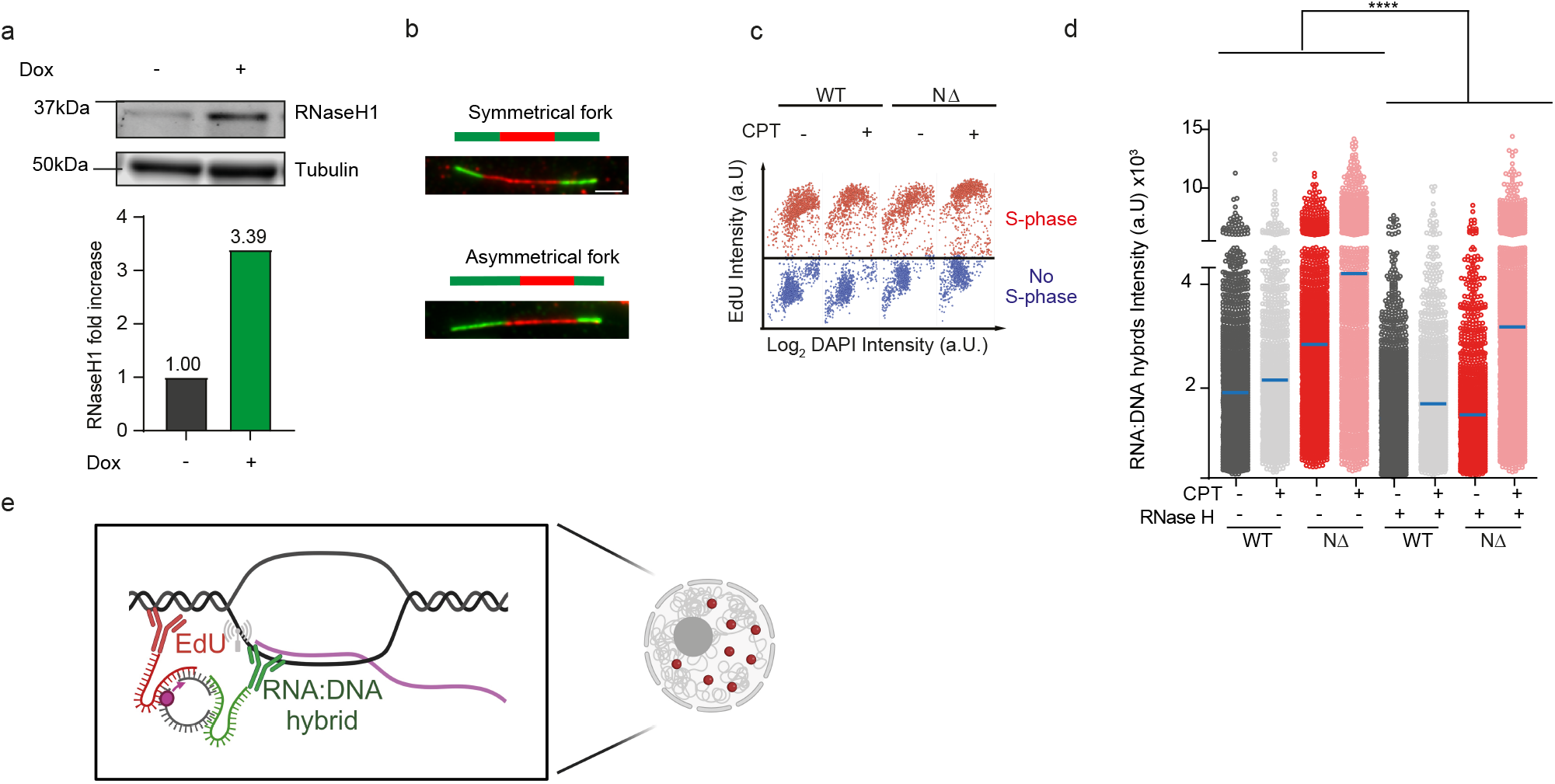
N-terminus region of SMARCAD1 is critical for R-loops resolution. **a.** Western blot analysis of Dox-inducible RNaseH1 over-expression. Uncropped membranes available in Source Data. **b.** Representative examples of sister fork asymmetry observed by sequentially labeling nascent replicating DNA in the indicated conditions. Scale bar, 5 μm. **c.** QIBC plot of EdU intensity over DAPI intensity upon CPT treatment separating EdU+ cells and EdU- cells in indicated cell lines and conditions. n(WT CPT) = 1700, n(NΔ CPT) = 1699. **d.** Quantification of high-content DNA-RNA hybrids intensity of the indicated conditions and treatments. n(WT UT) = 2780, n(WT CPT) = 3179, (WT RNaseH) = 1903 cell, n(WT CPT RNaseH) = 2042, n(NΔ UT) = 2389, n(NΔ CPT) = 2055, n(NΔ RNaseH) = 1903 cells, n(NΔ CPT RNaseH) = 2901 cells. ****=p≤0.0001, one-way analysis of variance Kruskal-Wallis test, followed by Dunn’s correction for multiple testing. Source numerical data available in Source Data. **e.** Cartoon of PLA principle between DNA-RNA hybrids and active replication forks. Figure was created with biorender.com.

**Extended Data Fig. 5.**
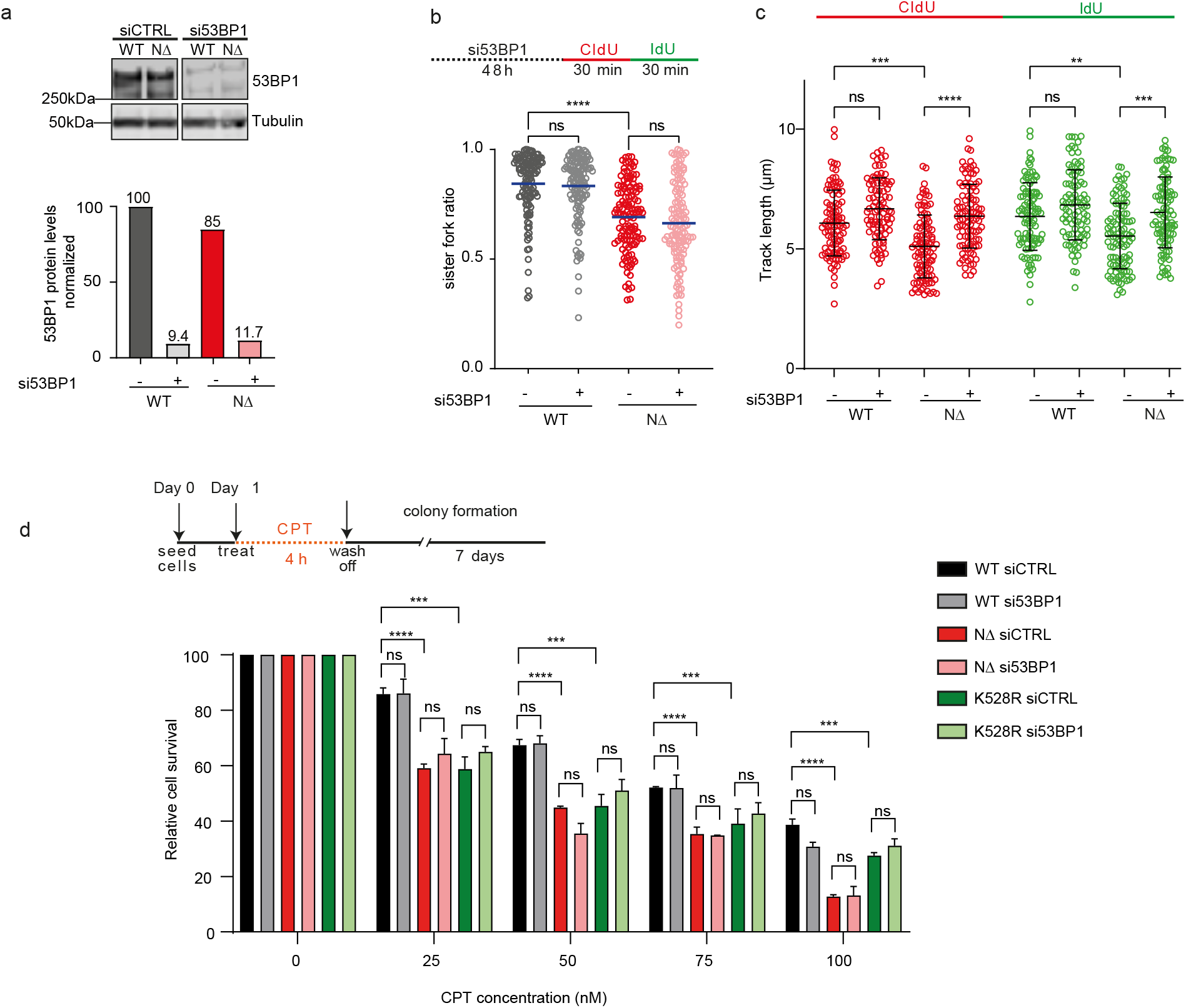
SMARCAD1 resolves R-loops independently from its PCNA homeostasis function. **a.** Western blot analysis of 53BP1 knock-down in indicated conditions. Uncropped membranes available in Source Data. **b.** Top: schematics of experimental set-up with knock-down of 53BP1 by siRNA and sequential labeling of nascent DNA. Bottom: quantification for sister fork ratio (ratio shorter/longer IdU tracks) of the indicated conditions. n(WT siCTRL) = 124, n(WT si53BP1) = 132, n(NΔ siCTRL) = 126, n(NΔ si53BP1) = 131 replicative sister forks. ns = non significant, ****=p≤0.0001, one-way analysis of variance Kruskal-Wallis test, followed by Dunn’s correction for multiple testing. Source numerical data available in Source Data. **c.** Quantification for track length for CldU (red) and IdU (green) of the indicated conditions. n(WT siCTRL) = 100, n(WT si53BP1) = 91, n(NΔ siCTRL) = 100, n(NΔ si53BP1) = 100 replicative sister forks. ns = non significant, **=p≤ 0.1, ***=p≤0.01, ****=p≤0.0001, one-way analysis of variance Kruskal-Wallis test, followed by Dunn’s correction for multiple testing. Source numerical data available in Source Data. **d.** Colony survival assay. Top: experimental set-up for CPT treatment and wash off and colony growth for 7 days. Mean survival in WT, NΔ-SMARCAD1 and K528-RSMARCAD1 in presence (siCTRL) or absence (si53BP1) of 53BP1 upon increasing concentrations of CPT treatment. Data is normalized to the 0 nM dose of the corresponding condition. Error bars represent standard deviation. n=2 independent experiments. ns = non significant, ***=p≤0.001, ****=p≤0.0001, ordinary two-way analysis of variance was used for multiple comparison. Numerical data available in Source Data.

**Extended Data Fig. 6.**
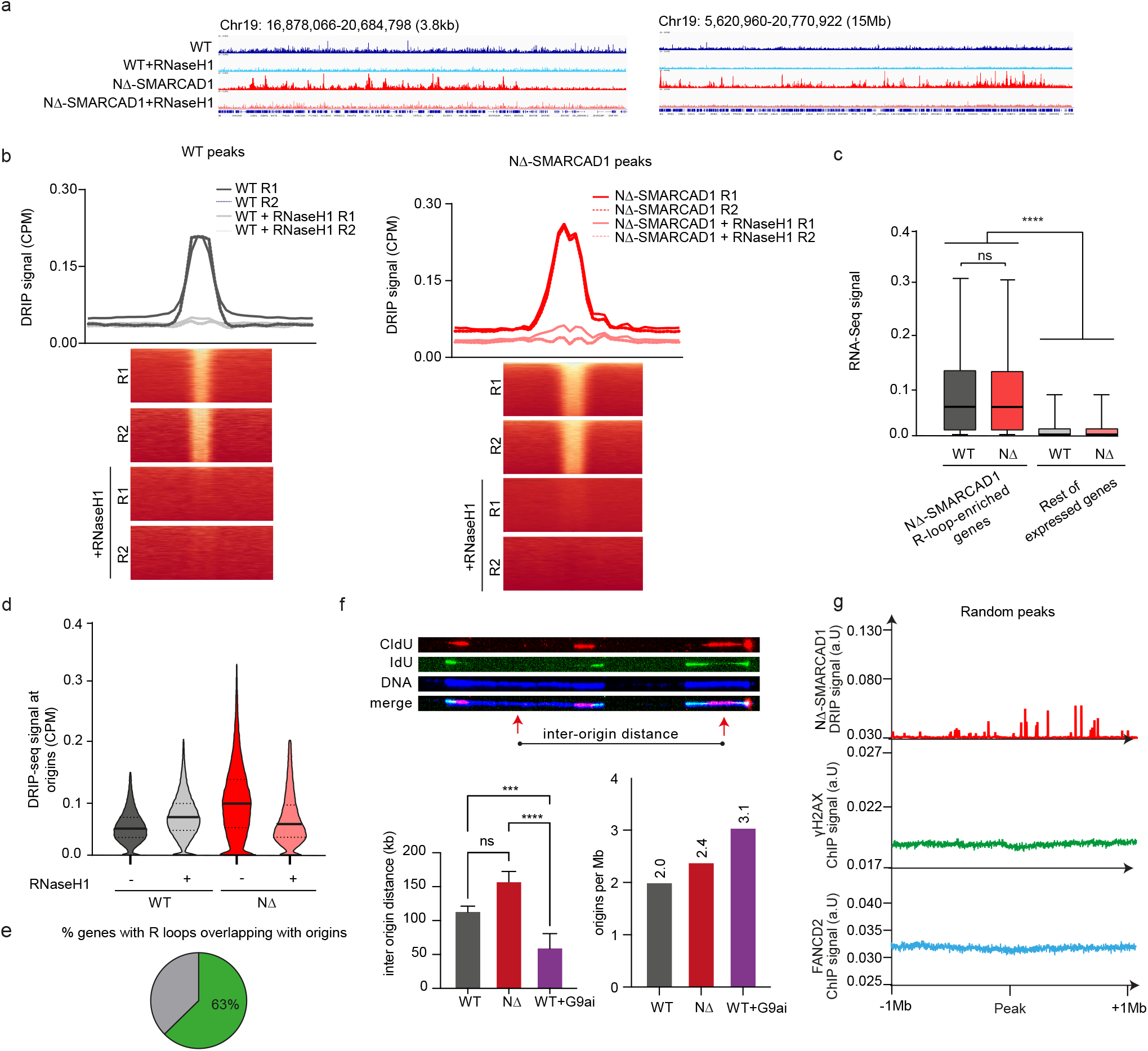
Loss of SMARCAD1 does not impair origin firing efficiency but leads to the accumulation of R-loops near replication origins, disrupting fork progression. **a.** Representative screenshots of genomic regions showing DRIP-seq signal from WT (blue) and NΔ-SMARCAD1 (red) cells treated (light) or not (dark) with RNaseH1 *in vitro*. **b.** Peak calling and heatmaps representation of WT (blue) and NΔ-SMARCAD1 (red) enriched peaks and their relative RNaseH1 in vitro treatment in two independent replicates. **c.** RNA-seq signal^42^ of genes around NΔ-SMARCAD1 R-loop-enriched sites compared to the rest of the genes in the indicated conditions. Numerical data available in Source Data. **d.** Violin plot representation of DRIP-Seq signal of the indicated conditions at origin of replication plotted by SNS-Seq^75^. ****=p ≤ 0.0001, one-way analysis of variance Kruskal-Wallis test, followed by Dunn’s correction for multiple testing. Numerical data available in Source Data. **e.** Distribution of R-loop-enriched genes (average coverage) overlapping with origins of replication. **f.** Top: Rep representative images of origins of replication frequency measurements. Bottom: Bar graph of inter-origin distance (left) and origin firing frequency (right) for WT, NΔ- SMARCAD1 and WT treated with 1 μM of G9a inhibitor for 2 h as in^90^, used as a positive control. ***=p≤0.001, ***=p≤0.0001, one-way analysis of variance Kruskal- Wallis test, followed by Dunn’s correction for multiple testing. Numerical data available in Source Data. **g.** Distribution of NΔ-SMARCAD1 DRIP-seq signal (red) and the ChIP-seq signals ± 1 Mb γH2AX (green) and FANCD2 (blue) from wild-type cells around random peaks.

**Extended Data Fig. 7.**
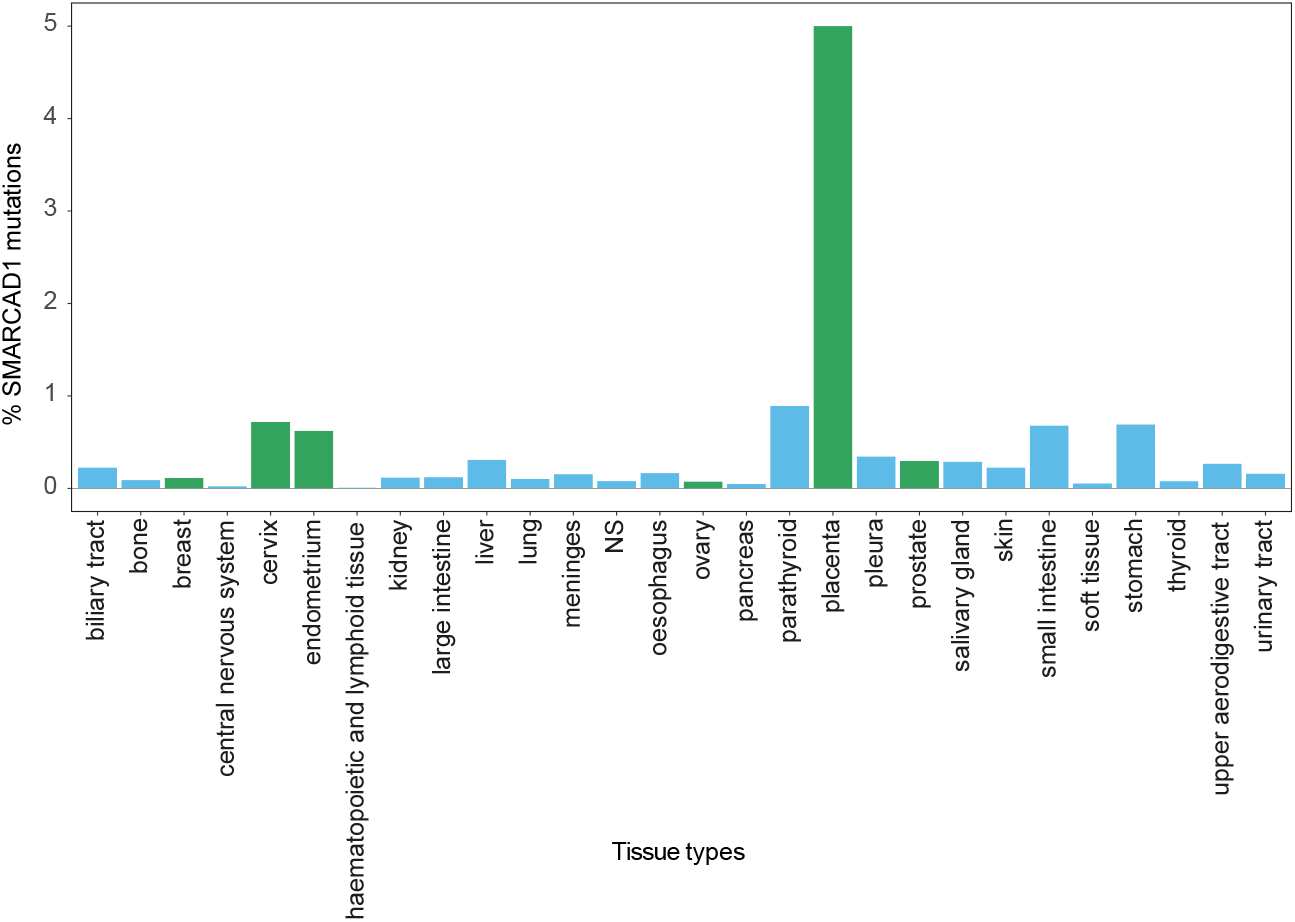
Mutational frequency of SMARCAD1 in different tissues. Analysis of percentage of mutation occurrence in different types of tissues in SMARCAD1.

**Supplementary Table 1.**

| **Cell line** | **Source** |
| --- | --- |
| MRC-5 sv40 immortalized (hereafter MRC5) | 42 |
| MRC-5 NΔ-SMARCAD1 | 42 |
| MRC-5 K528-SMARCAD1 | 42 |
| U2OS 2-6-3 | 59 |

**Supplementary Table 2.**

| **Compound and kits** | **Company (reference)** |
| --- | --- |
| 3-8% Criterion XT-Tri-Acetate Protein Gel | Bio-Rad (#3450131) |
| 4-12% Criterion XT-Tri-Acetate Protein Gel | Invitrogen (NP0335BOX) |
| 5'-chloro-2'-deoxyuridine (CldU) | Merck (C6891) |
| 5'-ethynyl-2'-deoxyuridine (EdU) | Sigma (D1306) |
| 5'-fluorescein label (6-FAM) | Integrated DNA Technologies |
| 5'-iodo-2'-deoxyuridine (IdU) | Merck (I7125) |
| 6% DNA retardation gel | ThermoFisher Scientific |
| Acetone | Honeywell (650501-1L) |
| Benzonase Nuclease HC, Purity >90% | Novagen (71205-3) |
| Biotin azide | Jena (CLK-1167-25) |
| Brilliant Blue R | Sigma (B0149) |
| BSA | Sigma (10735086001) |
| Chloroform | Sigma (C2432.2.5L) |
| Camptothecin (CPT) | Sigma (C9911) |
| DAPI | Sigma (E10187) |
| Doxycyclin | Sigma (D9891) |
| ECL reaction kit | Amersham (RNP2236) |
| FBS | Biowest |
| Formaldehyde 37% | Sigma (47608-1L-F) |
| GFP traps | Chromotek (GTA-200) |
| Glycogen | Thermo Scientific (R0561) |
| Ham's F10 | Invitrogen |
| Pen/Strep | Sigma-Aldrich |
| Phenol:Chloroform:Isoamyl acid 25:24:1 | ThermoFischer (10103971) |
| Duolink™ In Situ Detection Reagents Orange | Merck Millipore (DUO92007-100RXN) |
| Duolink™ In Situ PLA® Probe Anti-Rabbit PLUS | Merck Millipore (DUO92002-100RXN) |
| Duolink™ In Situ PLA® Probe Anti-Mouse MINUS | Merck Millipore (DUO92004-100RXN) |
| Prolong Gold Antifade mountant | Invitrogen (P36930) |
| Proteinase K | Roche (03115801001) |
| PVDF membranes | EMD Millipore (IPFL00010) |
| Quick Calf Intestinal Alkaline Phosphatase | New England BioLabs (M0525S) |
| RNAiMAX | ThermoFisher Scientific (56532) |
| RNaseH | New England BioLabs (M0297S) |
| 10X RNaseH reaction buffer | New England BioLabs (B0297S) |
| RNaseIII | Ambion (AM2290) |
| 10X RNaseIII buffer | Ambion (4019G) |
| Sepharose Cl-4B protecting column | GE Healthcare, Chicago |
| Sepharose Protein A binding column | GE Healthcare Chicago |
| SSC 20X | Sigma (SRE0068) |
| TAP300-Gold | Ted Pella |
| UNC0642 (G9a inhibitor) | MedChemExpress |

**Supplementary Table 3.**

| siRNA | Sequence | Company (reference) |
| --- | --- | --- |
| *53BP1* | L-003548-00-0020 | Dharmacon (7158) |
| Non Targeting Control pool | D-001810-10-20 | Dharmacon (1020) |
| *RNaseH1* | L-012595-01-0020 | Dharmacon (246243) |

**Supplementary Table 4.**

| Plasmid | Origin |
| --- | --- |
| pACEBAC1 | 34 |
| pFC53-mAIRN | 51 |
| mCherry-LacR-NLS-Stop | 58 |
| pLenti_CMV GFP puro | Luijsterburg lab |
| pLv_CMV-GFP-RNaseH1 | Luijsterburg lab |
| pCMV6-AC-GFP-SMARCAD1 | Gifted by van Attikum lab - <sup>49</sup> |
| pEBTet-BLAST-RNaseH1-myc/His | Gifted by Papamichos-Chronakis lab - <sup>25</sup> |

**Supplementary Table 5.**
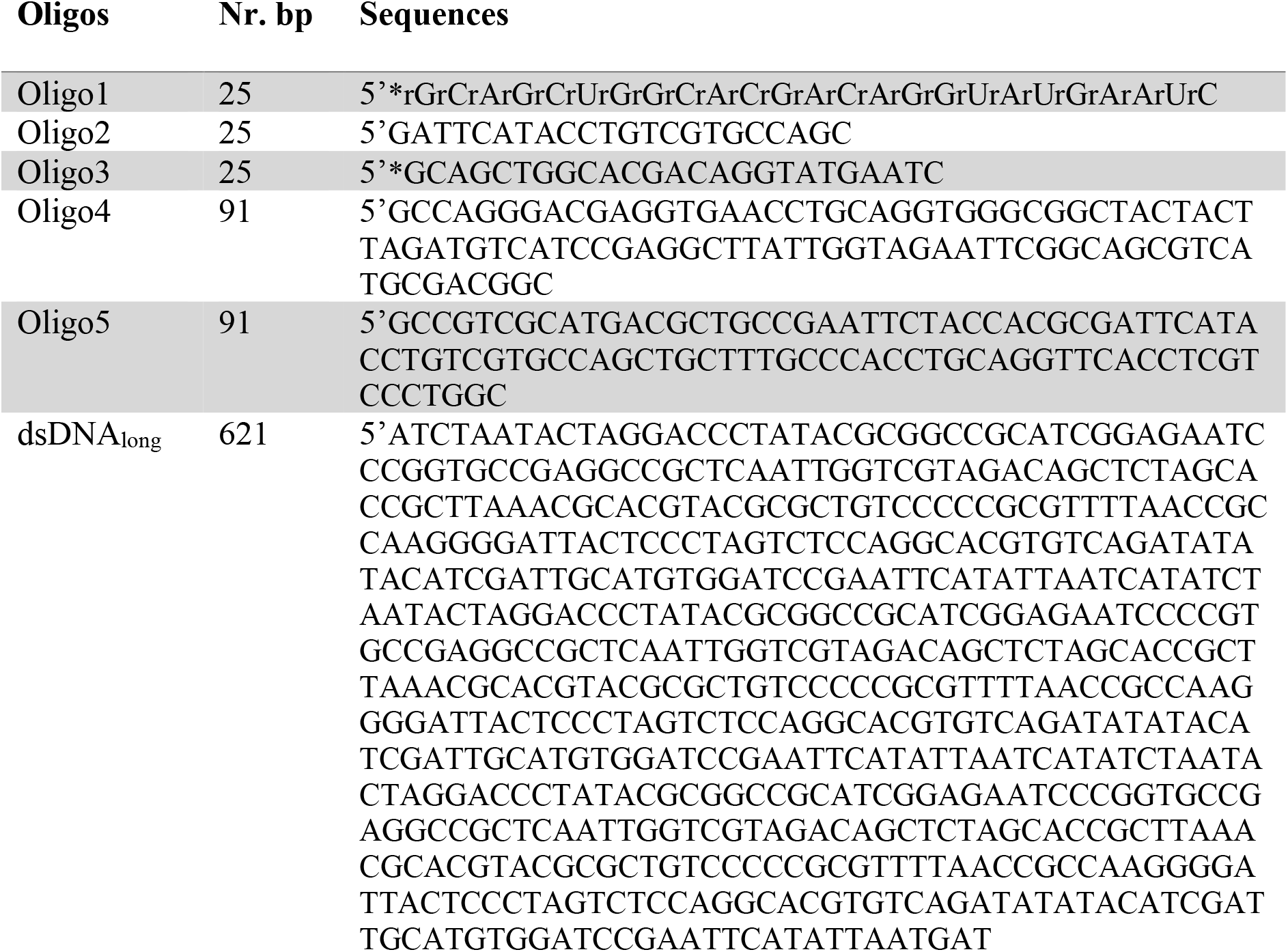

**Supplemental Table 6.**

| **Antibody** | **Species** | **Company (reference)** | **Use** |
| --- | --- | --- | --- |
| 53BP1 |  | Novus (NB100-304) | WB: 1:1000 |
| CldU (anti-BrdU clone BU1/75) | Rat | Abcam (ab6326) | IF: 1:200 |
| Autoanti-dsDNA |  | DSHB (AB_10805293) | WB: 1:1000 |
| IdU (anti-BrdU clone 44) | Mouse | Bectone Dickinson (#347580) | IF: 1:200 |
| HRP-conjugated secondary | Mouse | GE Healthcare (NA931V) | WB: 1:1000, PLA: 1:500 |
| HRP-conjugated secondary | Rabbit | GE Healthcare (NA934V) | WB: 1:1000 |
| Mouse IgG (H+L) Alexa 488 | Donkey | Jackson Immuno Research (715-545-150) | IF: 1:200 |
| Mouse IgG (H+L) Alexa 555 | Goat | ThermoFisher Scientific (A-21424) | IF: 1:200 |
| Mouse IgG(H+L) Alexa 647 | Goat | ThermoFisher Scientific (A-21235) | IF: 1:1000 |
| Mouse IgG(H+L) CF770 | Goat | Biotium (20077) | IF: 1:1000 |
| Rabbit IgG (H+L) Alexa 488 | Goat | ThermoFisher Scientific (A-11034) | IF: 1:1000 |
| Rabbit IgG (H+L) Alexa 555 | Goat | ThermoFisher Scientific (A-21229) | WB: 1:1000 |
| Rabbit IgG(H+L) Alexa 647 | Goat | ThermoFisher Scientific (A-21245) | IF: 1:1000 |
| Rabbit IgG(H+L) CF680 | Goat | Biotium (20067) | IF: 1:1000 |
| Rat IgG (H+L) Alexa 488 | Goat | ThermoFisher Scientific (A-21234) | IF: 1:1000 |
| RNaseH1 |  | Thermo Scientific (PA5-100003) | WB: 1:1000 |
| RNAPII-S2 | Rabbit | Abcam (ab5095) | WB: 1:1000 |
| S9.6 | Mouse | This study (GC lab) | High-content IF: 1:250<br>Artificial tethering IF: 1:1000 |
| S9.6 | Mouse | Hybridoma HB-8730 (AA lab) | DRIP |
| ssDNA | Mouse | Abcam (ab_10805144) | 1:50 |
| Tubulin | Mouse | Sigma (T5168) | IF: 1:1000 |

**Supplemental Table 7.**

| **Program** | **Use** |
| --- | --- |
| Adobe Illustrator 26.3.1 (64-bit) | Creation and assembling of figures |
| AlphaFold3 | Protein modeling in silico - <sup>55</sup> |
| CellProfiler | High-content imaging analysis |
| GraphPad Prism version 9.31 (471) | Data analysis and visualization |
| Gwyddion | AFM data visualization and analysis |
| IGV (version 2.7.7a) | Next generation sequencing analysis |
| Image Studio Lite Ver5.2 | Acquiring western blot scan |
| ImageJ 1.53g-Java 1.6.0_20 (64-bit) | Data analysis with personalized script |
| Mendeley (version 2.114.0) | Reference manager |
| Metasystem (version5) | High-content acquiring microscopy images and analysis |
| Microsoft® Excel® for Microsoft 365 MSO (Version 2208 Build 16.0.15601.20796) 32-bit | Data analysis |
| Microsoft® Word for Microsoft 365 MSO (Version 2208 Build 16.0.15601.20796) 32-bit | Text writing |
| PlotsOfData | Visualizing all datapoints - <sup>110</sup> |
| Spotfire (TIBCO) version 10.10.1 | QIBC data analysis and visualization |
| VolcanoseR | Visualizing volcano plots - <sup>111</sup> |
| Zen 2012 (blue edition) version 1.1.0.0 | Acquiring microscopy images |
| FASTQ Toolkit V.1.0.0 | DRIP-seq analysis |
| Rsubread V2.0.1 | DRIP-seq analysis |
| SAMtools V1.10 | DRIP-seq analysis |
| csaw V1.20.0 | DRIP-seq analysis |
| edgeR package (v3.20.9) | DRIP-seq analysis |
| ChIPpeakAnno V3.28.1 | DRIP-seq analysis |
| COSMIC v99 | Tumor analysis |

